# Quantitative Network Classification with CLaSSiNet Reveals Nanoscale Organizational Principles of the Membrane Skeleton

**DOI:** 10.64898/2025.12.20.695699

**Authors:** Yuan Tao, Ruobo Zhou

## Abstract

Quantitative analysis of nanoscale biological network architectures remains challenging in super-resolution fluorescence imaging, where cytoskeletal and network-like structures often appear as sparse localization clusters rather than continuous filaments. We introduce CLaSSiNet, a first-of-its-kind modular computational pipeline that integrates connectivity, 1D periodicity, and 2D regularity classifiers to automatically segment, classify, and map nanoscale network architectures with unprecedented classification resolution (∼256 nm) in single-molecule localization microscopy datasets. CLaSSiNet uniquely distinguishes four organizational states (1D periodic networks, 2D polygonal ordered networks, disordered networks, and non-network states) and is robust to experimental noise and compatible with both node- and link-based labeling strategies. Using CLaSSiNet, we achieve the first spatially resolved, quantitative mapping of organizational heterogeneity in the actin-spectrin membrane-associated periodic skeleton (MPS), a broadly conserved membrane scaffold in animal cells. We uncover previously unrecognized subcellular patterning, with ordered 1D and 2D networks enriched at cell edges and junctions, while non-network states dominate the cell body. CLaSSiNet further reveals mechanical coupling principles: actin stress fibers bias the formation and orientation of 1D periodic networks and enrich nearby non-network regions, suggesting coordination between spectrin lattices and contractile actin bundles. Comparative analysis across cell types shows that neurons favor 1D periodic architectures in neurites and 2D polygonal networks in somas, whereas U2OS and 3T3 cells adopt these architectures to a lesser extent, highlighting cell-type-specific tuning of spectrin network design. Together, these results establish CLaSSiNet as a generalizable platform for quantitative network-state mapping and provide new biological insights into how spectrin-based architectures adapt to local mechanical environments.

## Introduction

Super-resolution fluorescence microscopy, single-molecule localization microscopy (SMLM), has revolutionized our ability to visualize the nanoscale organization of cellular structures, revealing a rich diversity of network-like architectures that underline many fundamental biological processes^1–3^. These networks can often be abstracted into “nodes” (*e.g.*, protein complexes, filament junctions) physically connected by “links” (*e.g.*, cytoskeletal filaments, membrane tubules), and they span a wide range of dimensionalities, geometries, and degrees of structural regularity (**Figure 1A**). For instance, the membrane-associated periodic skeleton (MPS) in neurons predominantly forms a 1D periodic network in axons, where short actin filaments (nodes) are regularly spaced at 180-190 nm intervals and connected by spectrin tetramers (links of comparable length) (**Figure 1B**)^4^. In somas and dendrites, the same MPS components can give rise to a more variable 2D polygonal network, with less regular spacing and orientation (**Figure 1B**)^5,6^. The erythrocyte membrane skeleton represents a similar 2D polygonal network, composed of actin nodes and non-neuronal spectrin isoforms that generate average link lengths of 70-80 nm^7^. The sarcomere, the repeating unit of muscle fibers responsible for muscle contraction, exemplifies a 1D periodic contractile network, where actin and myosin filaments are precisely arranged between Z-discs spaced roughly 2.2 µm apart^8^. In contrast, networks such as the endoplasmic reticulum (ER) tubular system^9,10^ and tight junction strands^11–13^ in epithelial layers form more irregular polygonal architectures characterized by dynamic remodeling and substantial variability in link geometry, with link lengths often ranging from a few hundred nanometers to several micrometers. These diverse node-link networks play essential roles in supporting cellular architecture, enabling mechanical force transmission, organizing intracellular compartments, regulating molecular transport, and coordinating localized signaling events^13–16^. Despite their importance, there remains a lack of robust, automated computational tools to reliably identify, quantify, and classify such structurally diverse cellular networks, particularly in large-scale or noisy super-resolution imaging datasets^17^.

**Figure 1.**
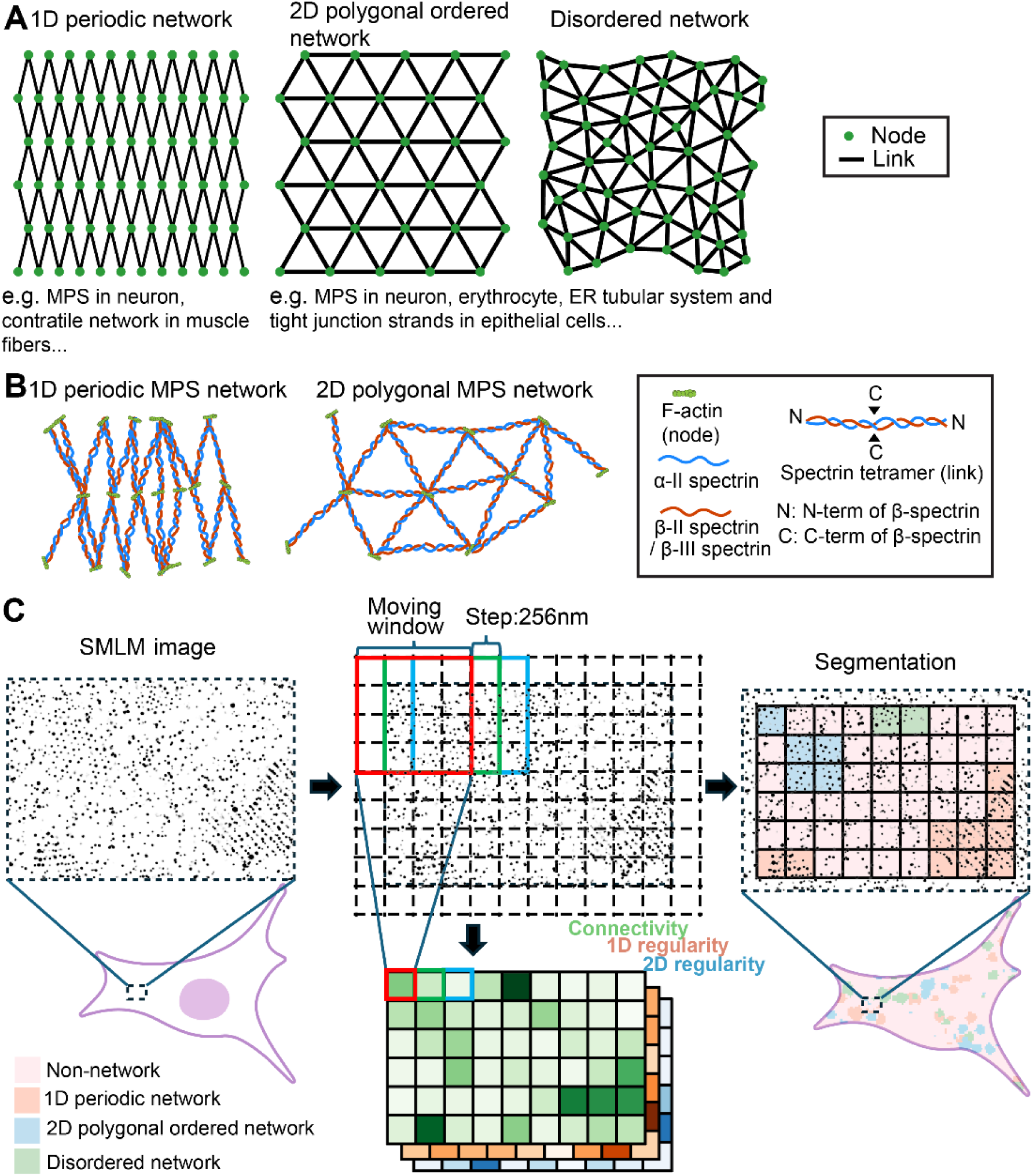
Example forms of network-like cellular architectures underlying fundamental biological processes and overview of the CLaSSiNet pipeline for their quantification and segmentation. **(A)** Schematics illustrating three distinct organizational states of network-like cellular architectures: one-dimensional (1D) periodic network, two-dimensional (2D) polygonal ordered network, and disordered network. **(B)** Schematics illustrating the organizations of 1D periodic MPS network and 2D polygonal MPS network. **(C)** Schematic overview of our developed CLaSSiNet pipeline for quantitative analysis and segmentation of super-resolution (SMLM) images of cellular networks. In each SMLM image, the cellular area containing signals is divided into a grid of 256 × 256 nm pixels. A 4 × 4–pixel moving window scans across the entire cellular area, and three classifier modules are applied within the moving window to compute likelihood scores corresponding to distinct organizational states. Each classifier module generates a heatmap representing one of three quantitative measures: connectivity (network-forming tendency), 1D regularity, and 2D regularity. The resulting heatmaps are then thresholded, binarized, and integrated to produce a segmentation map at 256 nm resolution, dividing the cellular area into four regions: 1D periodic, 2D polygonal ordered, disordered, and non-network regions.

Here, we present **C**lassifier of **S**uper-resolution **S**tructural **N**etworks (**CLaSSiNet**), a modular pipeline for the quantitative analysis of network-like cellular architectures from SMLM imaging data (**Figure 1C**). CLaSSiNet first identifies the cellular area in a SMLM image, then applies three classifier modules that segment the cellular area into four organizational sub-regions: regions primarily containing 1D periodic network, regions primarily containing 2D polygonal ordered network, regions primarily containing disordered network, and regions primarily containing non-network state. To enable high-resolution segmentation, the cellular area is divided into a grid with pixels of 256 × 256 nm. A moving window of 4 × 4 pixels (step size: 1 pixel) is then used as the analysis unit to scan across the entire cell area in the SMLM image. Each classifier module generates a heatmap representing one of three quantitative measures: connectivity (network-forming tendency), 1D periodicity, or 2D regularity, respectively. These heatmaps are subsequently thresholded, binarized, and integrated to generate a segmentation map with 256 nm resolution for each analyzed SMLM image.

Using the MPS, an actin-spectrin-based membrane skeleton present beneath the plasma membrane in many cell types^18–22^ and known to support essential cellular functions such as membrane protein anchoring, regulation of cellular mechanics, endocytosis, and membrane-associated signaling^23–26^, as a model system (**Figure 1B**), we applied CLaSSiNet to compare MPS organization in three different cell types: epithelial U2OS cells, fibroblast 3T3 cells, and neurons, uncovering previously unappreciated cell-type-specific organizational patterns. To further demonstrate the biological insights enabled by CLaSSiNet-based high-resolution network state segmentation, we analyzed how the MPS is organized around actin stress fibers, another major cytoskeletal structure formed at the adherent cell surface. We discovered that the likelihood of forming 1D periodic networks increases near stress fibers, and that the axis of MPS periodicity within these 1D-network-enriched regions is preferentially oriented perpendicular to the stress fibers. Together, these findings establish CLaSSiNet as a versatile and quantitative framework for dissecting nanoscale network organizational states across different cell types and subcellular contexts. By providing a systematic means to distinguish and compare diverse network architectures, CLaSSiNet enables the discovery of previously unknown structural principles that may govern how cytoskeletal and membrane-associated networks adapt to specific cellular environments. Beyond the MPS, CLaSSiNet offers a broadly applicable strategy for potentially probing the architecture of a wide variety of node-link networks in biology, opening avenues to uncover organizational rules that couple nanoscale patterning to cellular functions.

## Results

### Connectivity Classifier Module for Quantify the Network-forming tendency

In SMLM imaging, it is often not possible to directly visualize an entire biological network, including all nodes and their connecting links. Instead, immunofluorescence (IF) labeling typically reveals only the antigen positions where dye-conjugated antibodies bind. Therefore, reconstructing the node-link network of the biological networks from an SMLM image of a single cell requires a method that can infer the underlying connectivity based on the visualized nodes and quantifies local structural variations in network organization.

In this study, we used the actin–spectrin-based MPS as a model system for biological network analysis. The MPS is a ubiquitous and evolutionarily conserved cytoskeletal network underlying the plasma membrane of many cell types and is composed primarily of short actin filaments and actin-binding proteins, referred to as actin “nodes”, which are connected by spectrin tetramers that serve as “links”. The MPS can adopt either a one-dimensional (1D) periodic network, a two-dimensional (2D) polygonal ordered network, or disordered network, each covering certain portions of the plasma membrane (**Figure 1A, B**). IF-compatible antibodies have been developed to target the actin nodes, including antibodies that recognize short actin filaments or the N-termini of βII- or βIII-spectrin in cultured neurons^20^. These antibodies enable the acquisition of experimental SMLM images of actin node distributions (referred to as SMLM node images)

Because experimental SMLM node images do not provide ground truth network structures, which makes quantitative assessment of any network classifier impossible, we first aimed to establish a simulation method that generates node distributions with defined ground truth for each distinct organizational state of the MPS. Based on these experimental SMLM images obtained from neurons^20^, we developed a simulation framework that begins with a perfect 1D periodic or 2D polygonal ordered network (**Figure 1A**) and introduces a missed detection rate, a false-positive detection rate, and lateral positional deviations of nodes. These introduced perturbations were tuned such that the resulting simulated SMLM images reproduced the experimentally observed distributions of distances between nearest neighboring nodes for each organizational state. Using this approach, we simulated SMLM images of actin nodes arranged in three organizational states: (1) 1D periodic networks, (2) 2D polygonal ordered networks, and (3) randomly distributed nodes representing non-network regions (**Figure S1**). This simulation framework allows the generation of SMLM node images containing embedded “islands” of nodes arranged in any organizational state, such as 1D periodic, 2D polygonal ordered, on a background that can also adopt any combination of organizational states, providing a versatile and controlled dataset for assessing the performance of our classifier modules in segmenting SMLM images into sub-regions with distinct MPS organizational states.

Next, we developed our first classifier module, which reconstructs the node-link network of the MPS from SMLM node-only images and classifies the cellular area into network and non-network regions. To reconstruct the underlying network from SMLM node-only images, we considered two network reconstruction algorithms for the simulated SMLM node images (**Figure 2A; Figure S2A,B**): **(1)** Range-based network reconstruction: nodes within a predefined distance range around the average link length (*e.g.*, 190 ± 30 nm for MPS) were connected, and one of any intersecting links was randomly removed. **(2)** Delaunay-based network reconstruction: a Delaunay triangulation was generated from the node image, and links outside the predefined distance range (190 ± 30 nm) were removed except when only one link in a triangle was out of range and the other two links of the same triangle were within the range. To quantify the reconstructed network’s connectivity, defined as the degree of linkage per unit area, we used three measures based on the reconstructed network: (i) **link count**, number of links the reconstructed network contains per unit area, (ii) **triangle count**, number of triangles the reconstructed network contains per unit area (Triangles are defined as the smallest closed polygonal units in the reconstructed network, formed when three links connect neighboring nodes to enclose an area), and (iii) **triangle area sum**, total triangle area per unit area. Together, these produced six reconstruction–quantification pipelines (two reconstruction algorithms × three connectivity measures). Using the aforementioned moving-window approach to divide each SMLM image into grids (**Figure 1C**, **Figure 2A**), connectivity heatmaps can be generated across the entire cellular area in the SMLM image.

**Figure 2.**
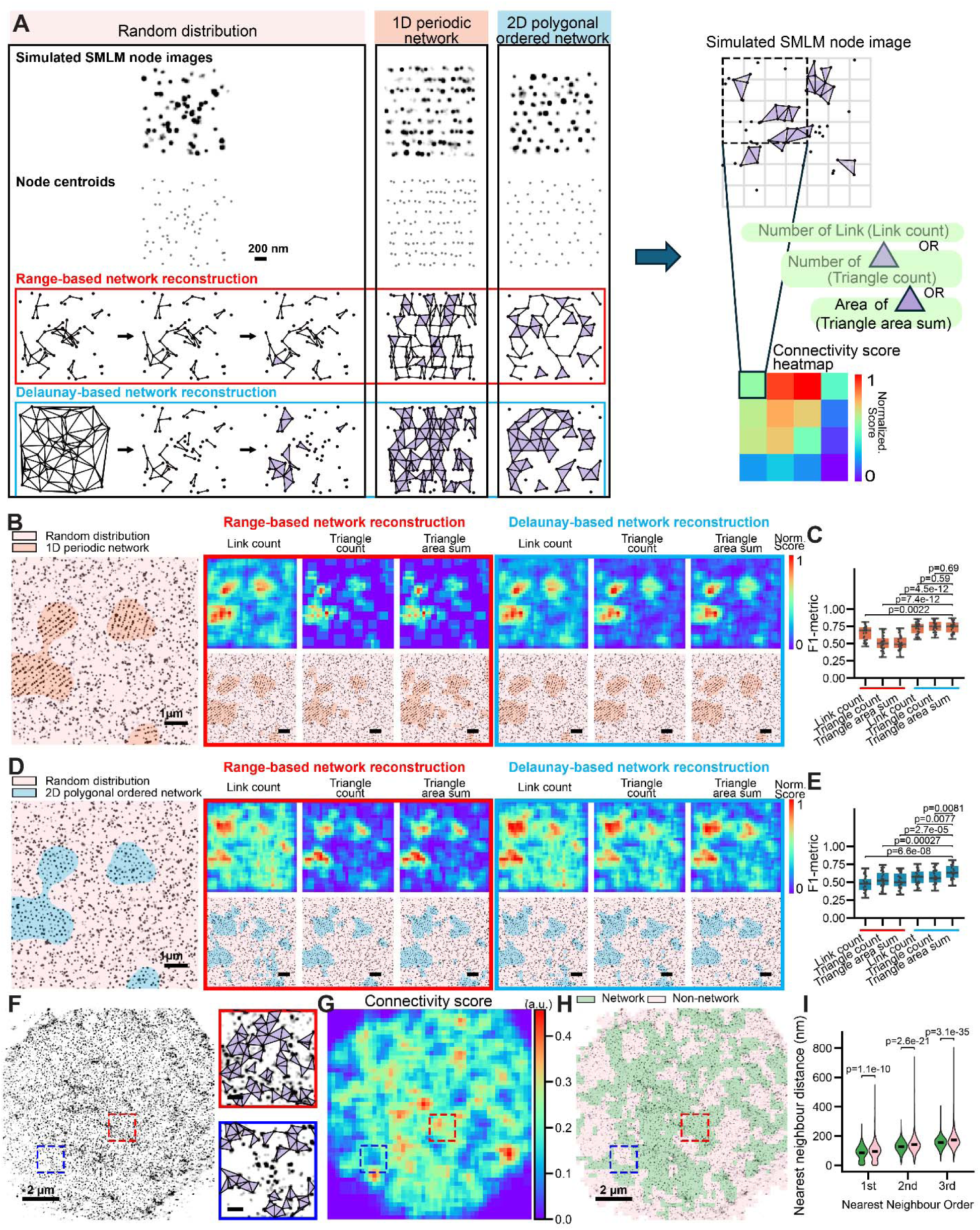
Connectivity Classifier Module for quantification of network-forming tendency and segmentation of network versus non-network cellular regions. **(A)** Top left: Representative simulated SMLM node images showing random, 1D periodic, and 2D polygonal distributions. Middle: Reconstructed networks generated from the three simulated images using range-based network reconstruction. Bottom left: Reconstructed networks obtained using Delaunay-based network reconstruction. The reconstruction steps are shown only for the simulated SMLM node image with random distribution. Right: Schematic illustrating three quantitative measures of connectivity (network-forming tendency), including link count, triangle count, and triangle area sum, and the generation of the corresponding connectivity heatmap. As shown in Figure 1c, each reconstructed network image was divided into 256 × 256 nm grids, and a 4 × 4-pixel moving window (step size: 1 pixel) was applied to compute the three measures and generate a connectivity heatmap.**(B)** Left: Representative simulated SMLM node image containing embedded islands of nodes arranged in a 1D periodic network, surrounded by randomly distributed nodes of equal density. Right: Connectivity heatmaps (top) and segmented images (bottom) generated for the simulated SMLM image on the left, using six reconstruction-quantification pipelines (two reconstruction algorithms × three connectivity measures). Scale bars: 1 µm. **(C)** Boxplots of F1-metrics quantifying the segmentation performance in distinguishing 1D periodic network regions from non-network (randomly distributed) regions. **(D,E)** Same as (B,C) but for simulated SMLM node images containing embedded islands of nodes arranged in a 2D polygonal network surrounded by randomly distributed nodes of equal density. **(F)** Left: Representative experimental SMLM image of βIII-spectrin (immunolabeled at its N-terminus). Right: Magnified views of the two boxed regions showing the reconstructed network. Scale bars: 2 µm. **(G, H)** Connectivity heatmap (G) and segmented image (H) generated for the SMLM image in (F). **(I)** Violin plots of the first, second, and third nearest-neighbor distances (NNDs), determined for network versus non-network regions. The center line indicate the median. *p*-values were calculated using a two-sided unpaired Student’s *t*-test.

To assess performance, we simulated SMLM node images containing embedded “islands” of nodes arranged in either a 1D periodic **(Figure 2B)** or 2D polygonal **(Figure 2D)** network, surrounded by nodes of equal density but random distribution. An effective reconstruction–quantification pipeline should sensitively detect the network-organized islands and distinguish them from the random background, which is expected to primarily consist of non-network nodes. Connectivity heatmaps generated from the six pipelines all successfully outlined the islands, albeit with varying levels of error. By applying thresholds for each connectivity measure (determined at a 5% false-positive rate, **Figure S2C,D**), we converted the heatmaps into binary segmentation maps of network versus non-network regions **(Figure 2B,D)**. Performance was quantified using the F1 metric^27^, which is the harmonic mean of precision and recall of a classification model in order to measure the model’s performance **(Figure2C,E)**. F1 metric analysis revealed that Delaunay-based network reconstruction combined with any of the three measures achieved higher F1 metrics than Range-based network reconstruction for distinguishing 1D periodic networks from random distributions. For 2D networks, the best performance was achieved with Delaunay-based network reconstruction combined with triangle area sum. Based on these results, we selected Delaunay-based network reconstruction with triangle area sum as the optimal pipeline, which we referred to as the Connectivity Classifier Module, as it most sensitively distinguished both 1D and 2D networked nodes from random (*i.e.* non-network) nodes.

To evaluate the sensitivity and robustness of the Connectivity Classifier Module to experimental noise, we applied it to simulated SMLM images of 1D and 2D networks with controlled perturbations. Three common noise sources were introduced: lateral positional deviations of node localization clusters, added false-positive nodes (e.g., fluorescent debris), and missed node detections. For each condition, we quantified connectivity scores and F1 metrics. Both measures decreased progressively with increasing noise, with the classifier showing high robustness to lateral deviations and false positives, whereas missed detections produced the strongest degradation in performance (**Figure S2E,F**). Notably, even for the least robust condition (*i.e.*, missed node detections), the F1 metric declined gradually and remained > 0.5 up to a 40–50% miss rate. Although an F1 metric of 0.5 usually represents the lower limit of acceptable performance, this result indicates that the classifier continues to extract meaningful network structure even under extreme signal loss.

Finally, we applied our Connectivity Classifier Module to experimental SMLM images of cultured hippocampal neurons immunostained for the N-terminus of βIII-spectrin, labeling the actin nodes (**Figure 2F**). The resulting binary segmentation maps showed that regions classified as “network” corresponded to areas with higher reconstructed link density (**Figure 2G,H**). To further characterize network organization, we performed nearest-neighbor distance (NND) analysis, which quantifies the distances between each node and its closest neighboring nodes within the network. Specifically, we calculated the first, second, and third NNDs for all nodes, where the first NND represents the distance to the closest neighboring node, the second NND to the next closest, and the third NND to the third closest neighbor. This analysis revealed that non-network regions exhibited significantly longer first, second, and third NNDs compared to network regions, reflecting the lower local node density and lack of organized connectivity in non-network areas, whereas network regions maintained shorter and more regular node spacing consistent with structured MPS organization (**Figure 2I**).

### 1D Network Classifier Module to Quantify the Regularity of 1D Periodic Networks

After segmenting SMLM node images into network and non-network regions using the Connectivity Classifier Module, we next sought to specifically identify and characterize 1D periodic networks, such as the 190-nm MPS organization observed in neuronal axons and dendrites^20,28^. To achieve this, we developed a two-dimensional Fast Fourier Transform (2D FFT)-based 1D Network Classifier Module that quantifies the degree of 1D periodic order in SMLM node images (**Figure 3A**).

**Figure 3:**
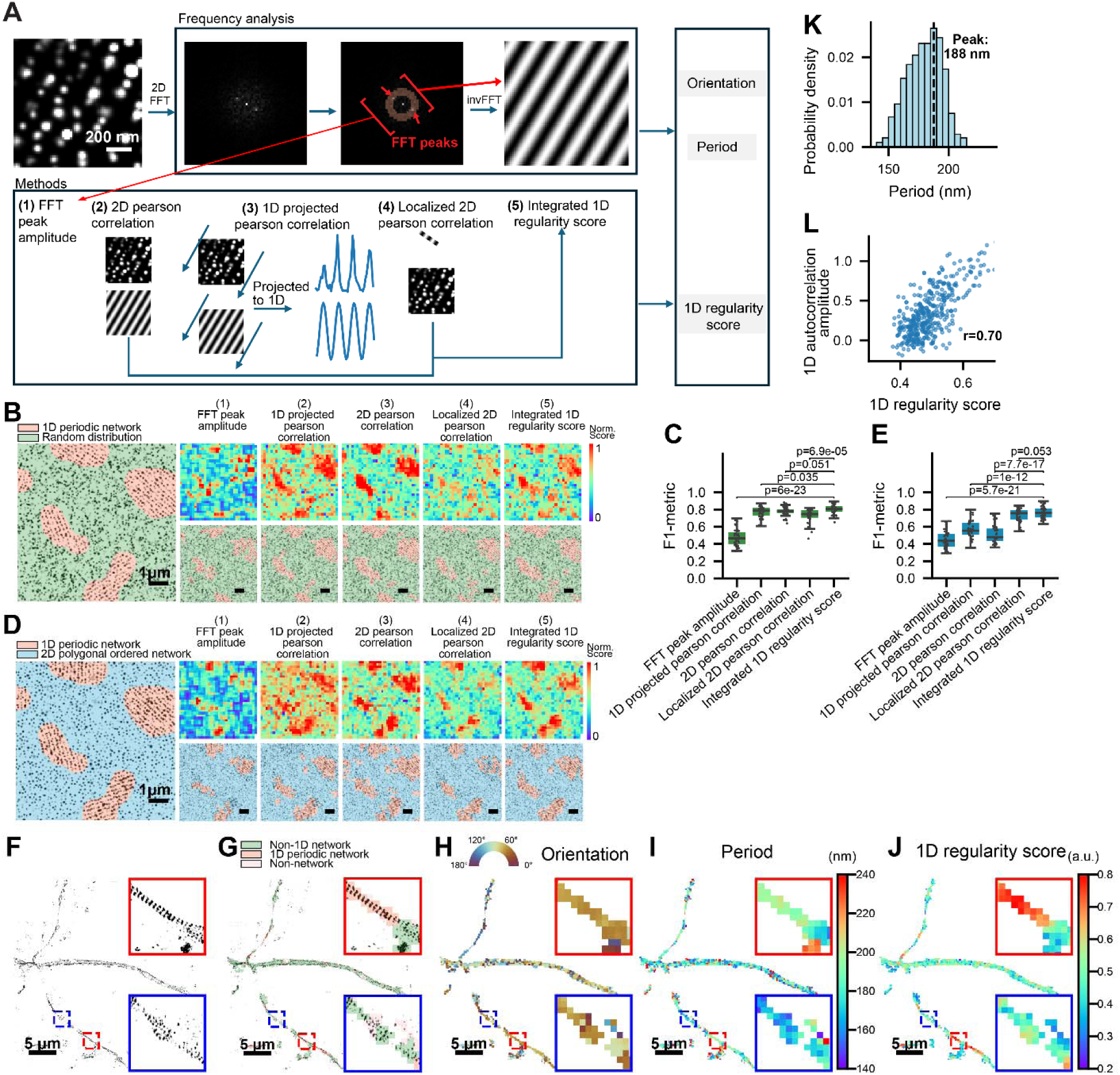
1D Network Classifier Module for quantification of 1D regularity and segmentation of 1D network versus non-1D network regions. **(A)** Schematic overview of the candidate quantification methods used to optimize the 1D Network Classifier Module. Within the moving window (1024 nm × 1024 nm), a two-dimensional Fast Fourier Transform (2D FFT) was performed to convert the spatial image into the frequency domain. In the resulting 2D frequency-domain image, the pair of peaks with the highest amplitudes within a donut-shaped region corresponding to a real-space period of 190 ± 30 nm were identified as the dominant frequency peaks. By retaining only these two peaks and removing all other frequency components, an inverse FFT (invFFT) was applied to generate a real-space grating image representing the best-fitting orientation and period of the potential 1D network within the moving window. Five quantitative 1D regularity scores were evaluated to assess the degree of 1D periodic order: (1) FFT peak amplitude, the amplitude of the two dominant frequency peaks; (2) 2D Pearson correlation, the correlation coefficient between the raw SMLM image and the invFFT-generated grating image; (3) 1D projected Pearson correlation, the correlation coefficient between 1D signal profiles obtained by projecting both images in (2) along the grating axis; (4) Localized 2D Pearson correlation, similar to (2) but computed using a cropped, elongated grating image to account for local 1D network regions; and (5) Integrated 1D regularity score, a combined metric that integrates measures (2) and (4) to capture both large 1D network islands spanning the full window and smaller local 1D regions. **(B)** Left: Representative simulated SMLM node image containing embedded islands of nodes arranged in a 1D periodic network, surrounded by randomly distributed nodes of equal density. Right: 1D regularity score heatmaps (top) and segmented images (bottom) generated for the simulated SMLM image on the left using the five 1D regularity scores described in (A). Scale bars: 11 µm. **(C)** Boxplots of F1-metrics quantifying segmentation performance in distinguishing 1D periodic network regions from random node distributions. **(D, E)** Same as (B, C), but for simulated SMLM node images containing embedded islands of nodes arranged in a 1D periodic network, surrounded by nodes forming a 2D polygonal network of equal density. Boxplots show the median and interquartile range (first and third quartiles); whiskers denote the minimum and maximum values excluding outliers. *p*-values were calculated using a two-sided unpaired Student’s *t*-test. **(F)** Experimental SMLM (STORM) image of βIII-spectrin in dendrites. Top right inset: magnified view of the boxed region showing high 1D regularity scores. **(G–J)** Segmented image (G), orientation heatmap (H), period heatmap (I), and 1D regularity score heatmap (J) generated for the SMLM image in (F). Scale bars: 2 µm. **(K)** Distribution of 1D periods obtained from the segmented 1D periodic regions identified in (J), showing a peak at ∼188 nm. **(L)** Scatter plot showing a strong positive correlation (*r* = 0.70) between 1D autocorrelation amplitudes determined by traditional 1D autocorrelation analysis and the 1D regularity scores determined by the 1D Network Classifier Module, calculated from the same set of experimental SMLM images.

This module used the aforementioned moving-window approach to divide each SMLM image into grids (**Figure 1C**). For each moving window (1024 nm × 1024 nm, with moving step 256 nm), a 2D FFT was performed to convert the spatial image into the frequency domain. In the resulting 2D frequency domain image, the pair of peaks with the highest amplitudes within a donut-shaped region corresponding to a real-space period of 190 ± 30 nm (the average period previously determined for 1D MPS) were identified as the dominant frequency peaks. By retaining only these two peaks and removing all other frequency components, an inverse FFT (invFFT) was applied to generate a real-space grating image representing the best-fitting orientation and period of the potential 1D network within the moving window. To quantify the degree of 1D periodic order, we calculated five candidate 1D regularity scores within the moving window: (1) FFT peak amplitude, the amplitude of the two dominant frequency peaks in the 2D frequency domain image; (2) 2D Pearson correlation, the Pearson correlation coefficient between the raw SMLM image and the invFFT-generated grating image; (3) 1D projected Pearson correlation, the Pearson correlation coefficient between 1D signal profiles obtained by projecting both images in (2) along the grating axis; (4) Localized 2D Pearson correlation, similar to (2), but the correlation is calculated between the raw SMLM image and a cropped, elongated grating image to account for cases where the 1D periodic network occupies only a subregion within the moving window; (5) Integrated 1D regularity score, a combined metric that mathematically integrates measures (2) and (4) to account for both large 1D network islands spanning the full window and smaller local 1D regions. As the moving window scans across the SMLM node image, each 1D regularity score generates a heatmap of the degree of 1D periodic order. In addition, the orientation and period associated with each window can be directly derived from the coordinates of the two dominant frequency peaks.

To compare the performance of these five 1D regularity scores, we simulated SMLM node images containing embedded islands with 1D periodic distributions, surrounded by either randomly distributed nodes or nodes forming 2D polygonal ordered networks of equal density (**Figure 3B,C).** Each 1D regularity score produced a corresponding heatmap of 1D periodic order. Because each heatmap pixel (256 × 256 nm) is smaller than the typical size of most 1D network islands, and the orientation and period values of adjacent pixels vary only gradually, we incorporated both parameters into a segmentation method based on a seed region growing algorithm (**Figure S3A,B**). Specifically, an initial thresholding corresponding to 1% false-positive rate (**Figure S3C**) was applied to the 1D regularity score heatmap to identify “seed” pixels with high 1D regularity score. The regions composed of these seed pixels were then expanded by including neighboring pixels that exceeded a second threshold corresponding to 5% false-positive rate (**Figure S3C**) and exhibited similar orientation and period values based on the orientation and period heatmaps. Using this refined segmentation method, the heatmaps were subsequently converted into binary maps distinguishing 1D network from non-1D network regions. F1-metric analysis demonstrated that the integrated 1D regularity score outperformed all other four scores, achieving the highest accuracy and sensitivity in distinguishing 1D islands from random or 2D-distributed nodes (**Figure 3D,E**).

To assess the sensitivity of the optimized 1D Network Classifier Module, simulated SMLM images with 1D periodic networks were perturbed with controlled experimental noise, including lateral localization deviations of nodes, false-positive nodes, and missed node detections. The 1D regularity score decreased progressively with increasing noise. Among the three noise types, the classifier was most robust to false-positive nodes and missed detections, whereas lateral localization deviations had the strongest impact and caused a more pronounced reduction in the F1 metric for identifying 1D regularity (**Figure S3D**).

We then applied the 1D Network Classifier Module to experimental SMLM images of hippocampal neuron dendrites immunostained for actin nodes (*i.e.*, the N-terminus of βIII-spectrin), which have been shown to exhibit mixtures of 1D, 2D, and non-network distributions in neurons. The resulting segmentation maps showed that regions classified as 1D network formed a subset of the broader “network” by the Connectivity Classifier Module, corresponding to areas with visually apparent 1D periodicity (**Figure 3F-J**). The distribution of measured periods across all identified 1D periodic network regions peaked at ∼188 nm (**Figure 3H, K**), consistent with previous reports^4,5^. Compared to the previously developed 1D autocorrelation analysis^20^, which requires manual selection of analyzed segments (segment length ≥ 2 μm to ensure reliable autocorrelation calculation) followed by projection of fluorescence signals onto the axis of the axonal or dendritic segment to calculate 1D autocorrelation amplitude as the measure for the degree of 1D periodic order, our module provides a much finer spatial resolution of 0.256 μm while simultaneously outputting period and orientation heatmaps. Notably, the 1D Network Classifier Module can identify 1D periodic subregions within neurite segments whose overall 1D autocorrelation amplitude would have classified the entire segment as non-1D network (**Figure S3E, F**), demonstrating more sensitive and robust detection of 1D networks compared to traditional autocorrelation analysis. Importantly, despite this higher resolution and sensitivity, our quantification correlated strongly with the autocorrelation method (r = 0.70, **Figure 3L**).

### 2D Network Classifier Module to Quantify the Regularity of 2D Networks

In the cellular regions classified as “network” by the Connectivity Classifier Module, we next aimed to further identify and quantify the 2D polygonal ordered network, in addition to the classification of 1D periodic network. We reasoned that, for any given node in a 2D polygonal network, its geometric features relative to its immediate neighboring nodes should resemble those of its neighbors more closely in an ordered network than in a disordered network. Therefore, to quantify the regularity (*i.e.* the degree of order) of 2D polygonal networks, we assessed the degree of similarity among neighboring nodes within the reconstructed node-link network (**Figure 4A**).

**Figure 4.**
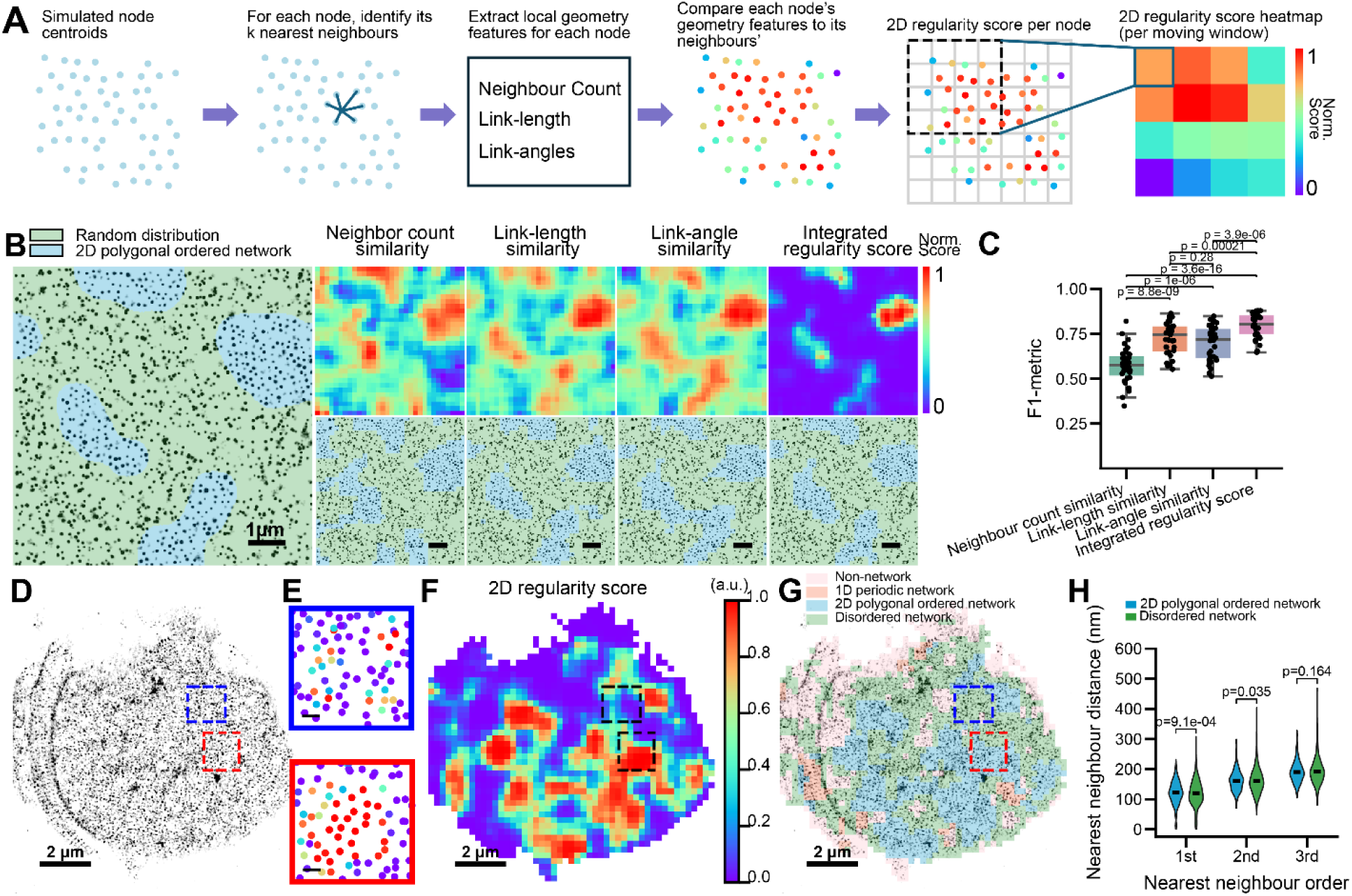
2D Network Classifier Module for quantification of 2D regularity and segmentation of 2D polygonal ordered network regions. **(A)** Schematic illustrating the steps of the 2D Network Classifier Module. Within the moving window (1024 nm × 1024 nm), the Connectivity Classifier Module was applied to reconstruct the node-link network. From the reconstructed network, three geometric features were calculated for each node: the number of connected neighboring nodes, the distribution of link lengths, and the distribution of link angles. The 2D regularity score for each node was defined as the inverse of the deviation of these geometric features from those of its immediate neighboring nodes, thereby quantifying the geometric self-consistency within local neighborhoods. The node-based 2D regularity scores were averaged across each moving window to generate 2D regularity score heatmaps. **(B)** Left: Representative simulated SMLM node image containing embedded islands of nodes arranged in a 2D polygonal network, surrounded by randomly distributed nodes of equal density. Right: Heatmaps (top) and segmented images (bottom) generated for the simulated SMLM image on the left, using the four candidate 2D regularity scoring methods. Scale bars: 1 µm. **(C)** Boxplots of F1-metrics quantifying the segmentation performance in distinguishing 2D polygonal network regions from random distributions. **(D)** Representative experimental SMLM image of βIII-spectrin (immunolabeled at its N-terminus). Scale bars: 2 µm. **(E)** Magnified views of the boxed region in (d) showing the reconstructed network with each node displaying the color-coded 2D regularity score. **(F, G)** 2D regularity score heatmap (F) and segmented image (G) generated for the SMLM image in (D). **(H)** Violin plots of the first, second, and third nearest-neighbor distances (NNDs), determined for 2D polygonal ordered network and disordered network regions. The center lines indicate the median. *p*-values were calculated using a two-sided unpaired Student’s *t*-test.

To quantify 2D polygonal network regularity, we applied the same moving-window approach described above (Figure 1c), partitioning the reconstructed node-link network from the SMLM node image into local grids, thereby enabling assessment of geometric consistency among nodes within each window. Specifically, for each moving window, we calculated four candidate 2D regularity scores based on three geometric features calculated for each node after network reconstruction from SMLM node images using the Connectivity Classifier Module: (1) **Neighbor count similarity**, the first geometric feature is the number of neighboring nodes directly connected to the given node. The 2D regularity score is defined as the inverse of the deviation in this neighbor count between a given node and its immediate neighbors; (2) **Link-length similarity**, the second geometric feature is the distribution of link lengths connecting the given node to its immediate neighbors. The corresponding 2D regularity score is defined as the inverse of the deviation in link-length distributions between a given node and its neighboring nodes; (3) **Link-angle similarity**, the third geometric feature is the distribution of angles formed between the links connected to the given node. The 2D regularity score is defined as the inverse of the deviation in angular distributions between a given node and its immediate neighbors; and (4) **Integrated 2D regularity score**, a composite 2D regularity score that simultaneously incorporates all three geometric features above. To determine the optimal combination of geometric features for distinguishing ordered from disordered networks, we employed Linear Discriminant Analysis (LDA). Using simulated ground-truth datasets containing 2D ordered and disordered networks, we trained an LDA model to identify the linear combination of the three individual 2D regularity scores that maximized the separation between ordered and disordered network classes. For each moving window, each 2D regularity score was calculated as the average value of all nodes within the moving window, generating a heatmap that represents the local degree of 2D regularity.

To compare the performance of these four 2D regularity scores, we simulated SMLM node images containing islands of nodes with 2D polygonal distributions embedded within backgrounds of either randomly distributed or 1D periodic nodes at identical node densities (**Figure 4B, Figure S4A**). Each 2D regularity score was applied to convert the simulated SMLM node image into a heatmap of 2D regularity, which was then thresholded to generate a binary segmented image. Quantitative comparison using the F1 metric showed that the intergrated 2D regularity score performed best in distinguishing 2D ordered networks from random node distributions (**Figure 4C**), whereas the link-angle similarity-based regularity score performed best in distinguishing 2D ordered networks from 1D periodic networks (**Figure S4A-C**). However, all four 2D regularity scores were generally less effective at distinguishing 2D networks from 1D periodic newtorks, than from random node distributions. Based on these results, we designate the integrated 2D regularity score, applied within the moving-window framework, as the 2D Network Classifier Module, which is most effective when applied after 1D periodic regions have been identified and removed by the 1D Network Classifier Module.

To assess the sensitivity and robustness of the optimized 2D Network Classifier Module, simulated SMLM images containing 2D polygonal networks were perturbed with controlled experimental noise, including lateral localization deviations, false-positive nodes, and missed detections, analogous to the tests performed for the Connectivity and 1D Network Classifier Modules. Both the 2D regularity score and the corresponding F1 metric decreased progressively with increasing noise levels (**Figure S4D**), demonstrating that the 2D classifier exhibits robustness comparable to that of the 1D Network Classifier Module.

Finally, we integrated all three modules into a complete analysis pipeline, referred to as **CLaSSiNet**, which segments an SMLM node image into four categories of cellular regions corresponding to distinct network organizational states: 1D periodic network, 2D polygonal ordered network, disordered network, and non-network. The workflow proceeds as follows: (1)Apply the Connectivity Classifier Module to segment the SMLM image into network and non-network regions; (2)Apply the 1D Network Classifier Module to identify 1D ordered networks within the network regions; (3)Apply the 2D Network Classifier Module to the remaining network regions to identify 2D ordered networks; and (4)Define the residual network regions as disordered networks.

Applying CLaSSiNet to our SMLM node images of the MPS network in neurons revealed distinct subregions corresponding to these four network organizational states (**Figure 4D-G**). We then performed nearest-neighbor distance (NND) analysis and found that 2D polygonal-ordered network regions exhibited NND values comparable to those of disordered network regions (**Figure 4H**).

### Performance of Classifier Modules on SMLM Images Labeling Link-Midpoints Instead of Nodes

As immunofluorescence-based SMLM often visualizes antigens as point localizations on node-link networks, we next examined how our three classifier modules would perform when the labeled antigen is located not at the nodes but at the midpoints of links. Using the MPS network as a model system, we compared antibodies recognizing the N-terminus of βII- or βIII-spectrin, which localize to actin nodes, with antibodies recognizing the C-terminus of βII-spectrin, which localizes to the midpoints of spectrin tetramers. In a 1D periodic network, SMLM images of link-midpoints are expected to exhibit periodic patterns similar to SMLM node images. By contrast, in a 2D polygonal network, link-midpoint distributions differ from node distributions (**Figure 5A**). This raised the question of whether our three classifier modules optimized on node images would also apply to link-midpoint images, and whether the two representations would yield similar classification and segmentation results.

**Figure 5.**
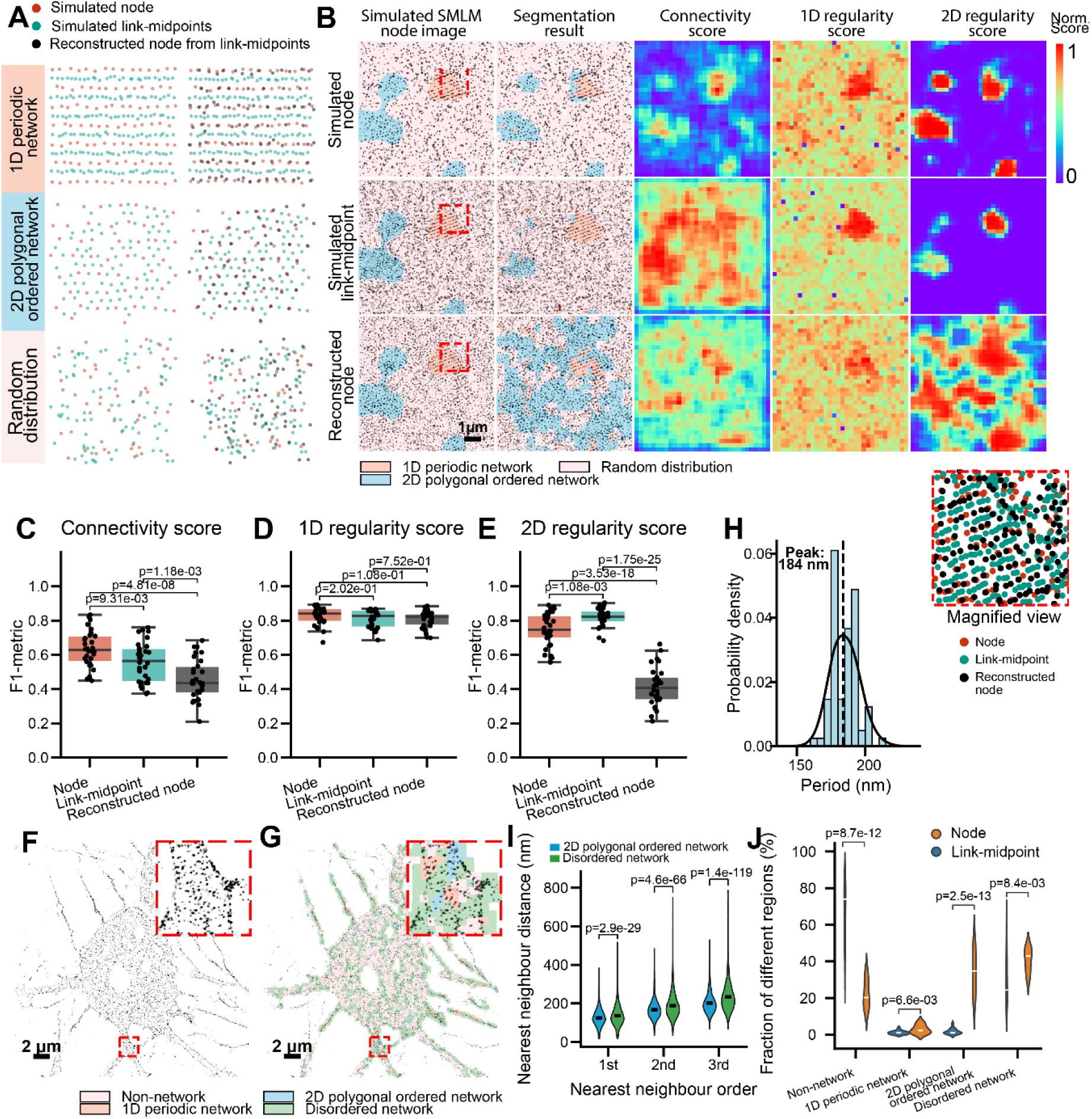
Performance of classifier modules on SMLM images labeled at link midpoints. **(A)** Simulated SMLM images illustrating node versus link-midpoint labeling for the same underlying networks: 1D periodic network (top row), 2D polygonal ordered network (middle row), and random distribution (bottom row). Only the centroids of molecular clusters in each SMLM image are shown. **(B)** Left: Representative simulated SMLM image containing embedded network islands composed of molecular clusters arranged in either a 1D periodic or a 2D polygonal ordered distribution, surrounded by randomly distributed clusters. Middle and Right: Corresponding segmented maps (middle) as well as connectivity, 1D regularity, and 2D regularity score heatmaps (right) generated using CLaSSiNet. A magnetified view of the red boxed region is shown at the bottom right corner. Scale bars: 1 µm. **(C–E)** Boxplots of F1 metrics quantifying segmentation performance for the three simulated SMLM image sets shown in (B): Connectivity Classifier Module (C), 1D Network Classifier Module (D), and 2D Network Classifier Module (E). **(F)** Representative experimental SMLM (STORM) image of βII-spectrin immunolabeled at the C-terminus (link-midpoint labeling) in a cultured neuron. **(G)** Segmented image corresponding to the SMLM image in (F). **(H)** Distribution of 1D periodicities obtained from the segmented 1D periodic regions in (F), showing a peak at ∼184 nm. Scale bars: 2 µm. **(I)** Violin plots of the first, second, and third nearest-neighbor distances (NNDs) measured from the segmented 2D polygonal ordered network and disordered network regions, with the median shown as the center line. **(J)** Violin plots showing the fractions of cellular area in STORM images occupied by the four distinct MPS structural states. Fractions were calculated separately for neuronal somas labeled at βII-spectrin C-termini (link-midpoints) or βIII-spectrin N-termini (nodes). The center lines indicate the median. *p*-values were calculated using a two-sided unpaired Student’s *t*-test.

To quantitatively test this, we generated three sets of simulated SMLM images containing embedded network “islands,” each composed of molecular clusters arranged in either a 1D periodic or a 2D polygonal ordered pattern, surrounded by randomly distributed clusters: (1) **Set 1**, SMLM node images with embedded islands exhibiting either 1D periodic or 2D polygonal ordered distributions; (2) **Set 2**, the corresponding link-midpoint version of Set 1, in which the same underlying networks were labeled at link midpoints instead of nodes; and (3) **Set 3**, predicted SMLM node images computationally inferred from Set 2 using a Voronoi tessellation-based algorithm to estimate node locations from link-midpoint distributions (**Figure S5**). Set 1 served as the ground truth for evaluating the predicted node images (Set 3). We then applied the three classifier modules to all three datasets to generate connectivity, 1D regularity, and 2D regularity score heatmaps and their corresponding segmentation results (**Figure 5B**). Quantitative comparisons using the F1 metric revealed that directly applying the three classifier modules to SMLM link-midpoint images (Set 2) yielded better performance than predicting node positions from the same data and then analyzing the inferred node images (Set 3). Furthermore, while segmentation results from SMLM link-midpoint images (Set 2) showed comparable accuracy to those from node images (Set 1) for the 1D Network Classifier Module, they exhibited slightly reduced performance for the Connectivity Classifier Module and slightly better performance for the 2D Network Classifier Module (**Figure 5C-E**).

Together, these results demonstrate that our three classifier modules can robustly quantify and classify SMLM images of node-link networks regardless of whether the fluorescent label marks nodes or link midpoints, and that predicting node distributions from midpoint data is less effective.

We further applied the CLaSSiNet pipeline to experimental SMLM images labeling spectrin link midpoints (*i.e.*, C-terminus of βII-spectrin) in the somas of cultured neurons and successfully segmented the networks into 1D periodic, 2D polygonal ordered, disordered, and non-network regions (**Figure 5F,G**). The distribution of periods identified in the segmented 1D periodic network regions (**Figure 5H**) was comparable to those obtained from βIII-spectrin N-terminus-labeled (i.e., node-labeled) SMLM images (**Figures 3K, 5H**). The NND analysis of βII-spectrin C-terminus–labeled SMLM images showed that 2D polygonal-ordered network regions exhibited NND values comparable to those of disordered network regions (**Figure 5I**), consistent with observations from βIII-spectrin N-terminus–labeled (*i.e.*, node-labeled) SMLM images (**Figure 4H**). However, when comparing the fractions of cellular regions exhibiting different MPS organizational states, βII-spectrin C-terminus labeling yielded smaller area fractions of 1D periodic, 2D polygonal-ordered, and disordered networks (**Figure 5J**). This likely reflects the spectrin isoform composition in neurons: βIII-spectrin is substantially more abundant than βII-spectrin in somas and dendrites of neurons, and thus βIII-spectrin labeling provides a more complete representation of the MPS network in these regions than βII-spectrin labeling.

### Application of CLaSSiNet Reveals Distinct Subcellular MPS Organizations in Epithelial and Fibroblast Cells

Having developed the integrated pipeline CLaSSiNet which incorporates three classifier modules to quantify and segment SMLM images of either nodes or link-midpoints in a node-link network, we next applied it to experimental SMLM data obtained from non-neuronal cells. CLaSSiNet classifies SMLM image regions into four distinct MPS organizational states, including 1D periodic networks, 2D polygonal ordered networks, disordered networks, and non-network regions. Recent studies have revealed that the spectrin-based membrane skeleton (herein also referred to as the MPS), previously well characterized in neurons and erythrocytes using SMLM^19,2820^, is also present in mouse embryonic fibroblasts (MEFs)^22,29^. In MEFs, the MPS was found depleted at sites occupied by actin stress fibers, the contractile bundles of actin filaments and myosin proteins that anchor the cell to the extracellular matrix through focal adhesions and enable cell attachment and spreading. The 1D MPS network, resembling those found in neuronal axons and dendrites, were observed between stress fibers at the adhesive surface of MEFs using expansion microscopy (ExM)^29^. However, whether similar MPS organizations exist in the epithelial cell type remains unclear.

To address this question, we first immunostained an epithelial cell line, U2OS, for two structural components of the MPS: βII-spectrin (a ubiquitous β-spectrin isoform expressed in most non-erythrocyte cell types) and F-actin^4^. Confocal imaging revealed that βII-spectrin displayed a heterogeneous distribution at the adhesive surface, similar to that previously reported in MEFs^29^. F-actin staining showed mutually exclusive pattern relative to βII-spectrin, further supporting their spatial segregation. To compare MPS organization across different subcellular regions, we segmented the cell area into four distinct zones: nucleus, cell body, cell–cell junction, and cell edge (**Figure 6B,C**). Quantification of the average fluorescence intensities of βII-spectrin and F-actin across these subcellular zones revealed that βII-spectrin was significantly enriched at cell edges and cell–cell junctions, whereas F-actin levels remained relatively uniform across all three non-nuclear zones (**Figure 6D**). This observation raised the possibility that distinct MPS organizational states may occupy different proportions of the three subcellular zones. To test this, we acquired SMLM images of βII-spectrin using Stochastic Optical Reconstruction Microscopy (STORM), a SMLM imaging method^1–3^, and applied CLaSSiNet for quantitative segmentation at a spatial resolution of 256 nm. Indeed, MPS at cell edges and cell-cell junctions contained higher fractions of 1D periodic and 2D polygonal ordered regions, whereas the cell body contained more non-network regions and dominated by disordered networks (**Figure 6E–H**). We further quantified the physical properties of these organizational states: 1D periodic network regions exhibited larger periods at cell edges and cell–cell junctions compared to the cell body, while regions classified as 2D polygonal ordered had more comparable NND distributions and means across the three subcellular zones (**Figure 6I, J**).

**Figure 6.**
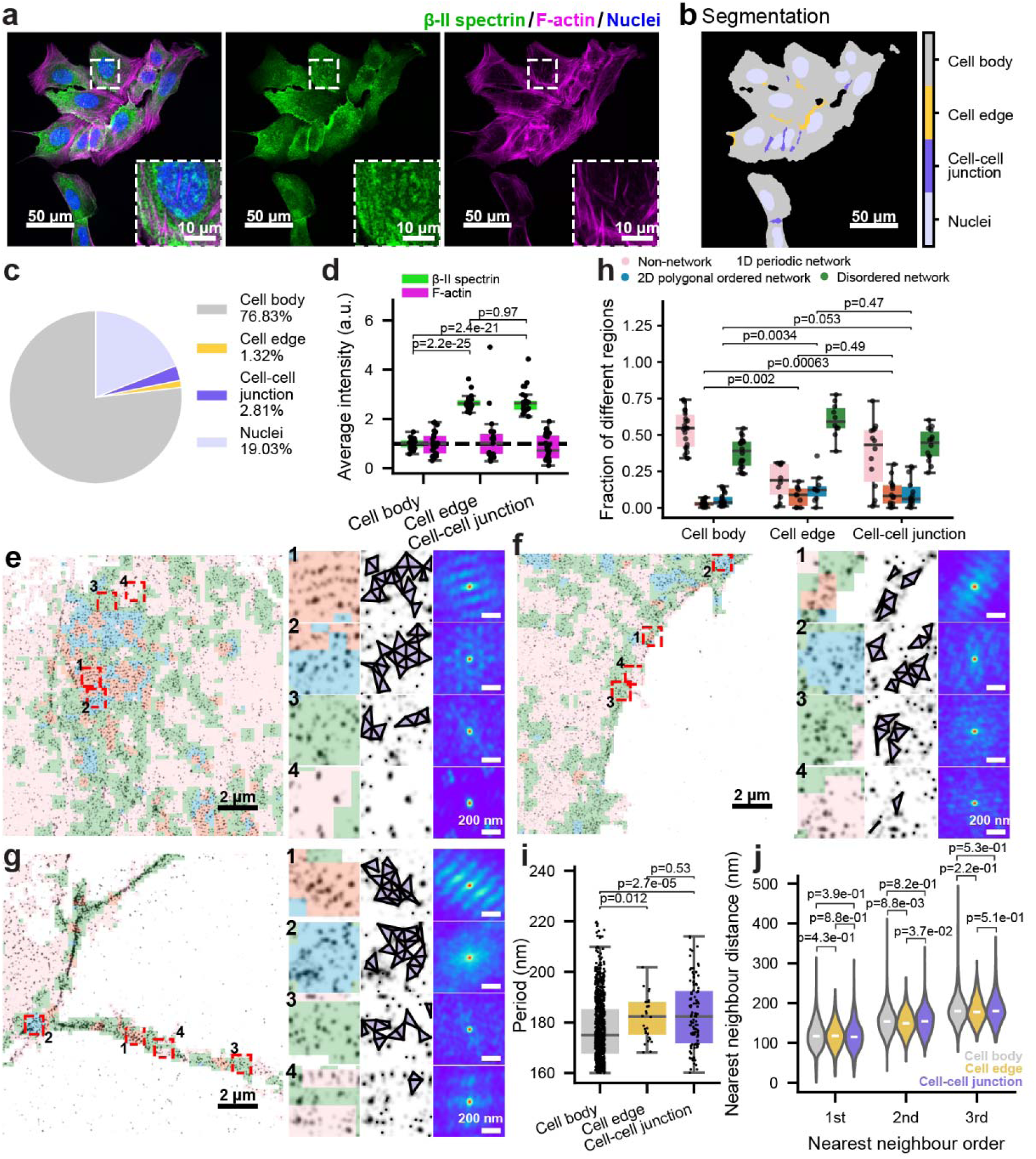
CLaSSiNet reveals distinct MPS organizations across three subcellular zones in epithelial cells. **(A)** Representative three-color confocal images of U2OS cells co-stained for nuclei (blue, Hoechst), βII-spectrin (green, βII-spectrin antibody), and F-actin (magenta, phalloidin). Scale bar: 50 µm. Inset: Magnified view of the boxed region. Scale bar: 10 µm. **(B)** Segmented image corresponding to (A), showing four subcellular zones: nucleus, cell body, cell edge, and cell–cell junction. Scale bar: 50 µm. **(C)** Quantified area fractions of the four subcellular zones. **(D)** Boxplots showing average fluorescence intensities of βII-spectrin and F-actin across the three non-nuclear subcellular zones (cell body, cell edge, and cell–cell junction). **(E–G)** Left: Representative SMLM (STORM) images of βII-spectrin in three subcellular zones, cell body (E), cell edge (F), and cell–cell junction (G), overlaid with CLaSSiNet segmentation into four MPS organizational states: 1D periodic network, 2D polygonal ordered network, disordered network, and non-network regions. Scale bar: 2 µm. Middle: Magnified views of boxed regions enriched in each of the four organizational states. Right: Reconstructed node-link networks and corresponding 2D autocorrelation maps confirming the expected MPS organizational patterns. Scale bar: 200 nm. **(H)** Boxplots showing the area fractions of the four MPS organizational states across the three subcellular zones. **(I)** Boxplots of 1D periodicity periods measured from 1D periodic network regions across the three subcellular zones. Boxplots show the median and interquartile range (first and third quartiles); whiskers denote the minimum and maximum values excluding outliers. Boxplots show the median and interquartile range (first and third quartiles); whiskers denote the minimum and maximum values excluding outliers. **(J**) Violin plots comparing the first, second, and third nearest-neighbor distance (NND) distributions of 2D polygonal ordered network regions across the three subcellular zones, with the median shown as the center line. *p*-values were calculated using a two-sided unpaired Student’s *t*-test.

We next performed a parallel analysis in fibroblast 3T3 cells. Although confocal imaging showed enrichment of spectrin and F-actin at cell edges and junctions, SMLM imaging combined with our **CLaSSiNet** analysis revealed results similar to those observed in U2OS cells, expect that 1D periodic and 2D polygonal ordered regions have a higher fraction only at cell edges but not at cell-cell junctions (**Figure S6**), likely because that fibroblast cell has less defined cell-cell junctions than epithelial cells.

### Stress Fiber Proximity Influences Local MPS Organization

Recent studies indicate that the MPS and actin stress fibers are largely non-overlapping at the cell surface (**Figure 7A**), leading us to hypothesize that local MPS organization, whether 1D periodic, 2D polygonal ordered, disordered network, or non-network, may depend on distance to nearby stress fibers. To test this, we peformed STORM imaging to obtain SMLM images of the βII-spectrin C-terminus (located at the midpoint of a spectrin tetramer, *i.e.*, link-midpoints) in U2OS cells, while simultaneously visualizing actin stress fibers using conventional fluorescence microscopy (**Figure 7B**). Applying CLaSSiNet to segment the SMLM images into four organizational states allowed us to quantify how the fraction of regions exhibiting each state varies as a function of distance to the nearest stress fiber. We found that non-network and 1D periodic network regions were enriched near stress fibers, whereas 2D polygonal ordered network and disordered network regions were depleted near them (**Figure 7E**), suggesting that proximity to stress fibers promotes 1D periodic organization. We next examined whether 1D periodic regions exhibit a preferred orientation relative to stress fibers. Orientation measurements from our 1D classifier module revealed a sharp peak at ∼90° relative to stress fiber orientation, indicating that the axis of MPS periodicity (*i.e.*, orientation of the periodic pattern) is preferentially aligned perpendicular to the nearest stress fiber (**Figure 7F**).

**Figure 7.**
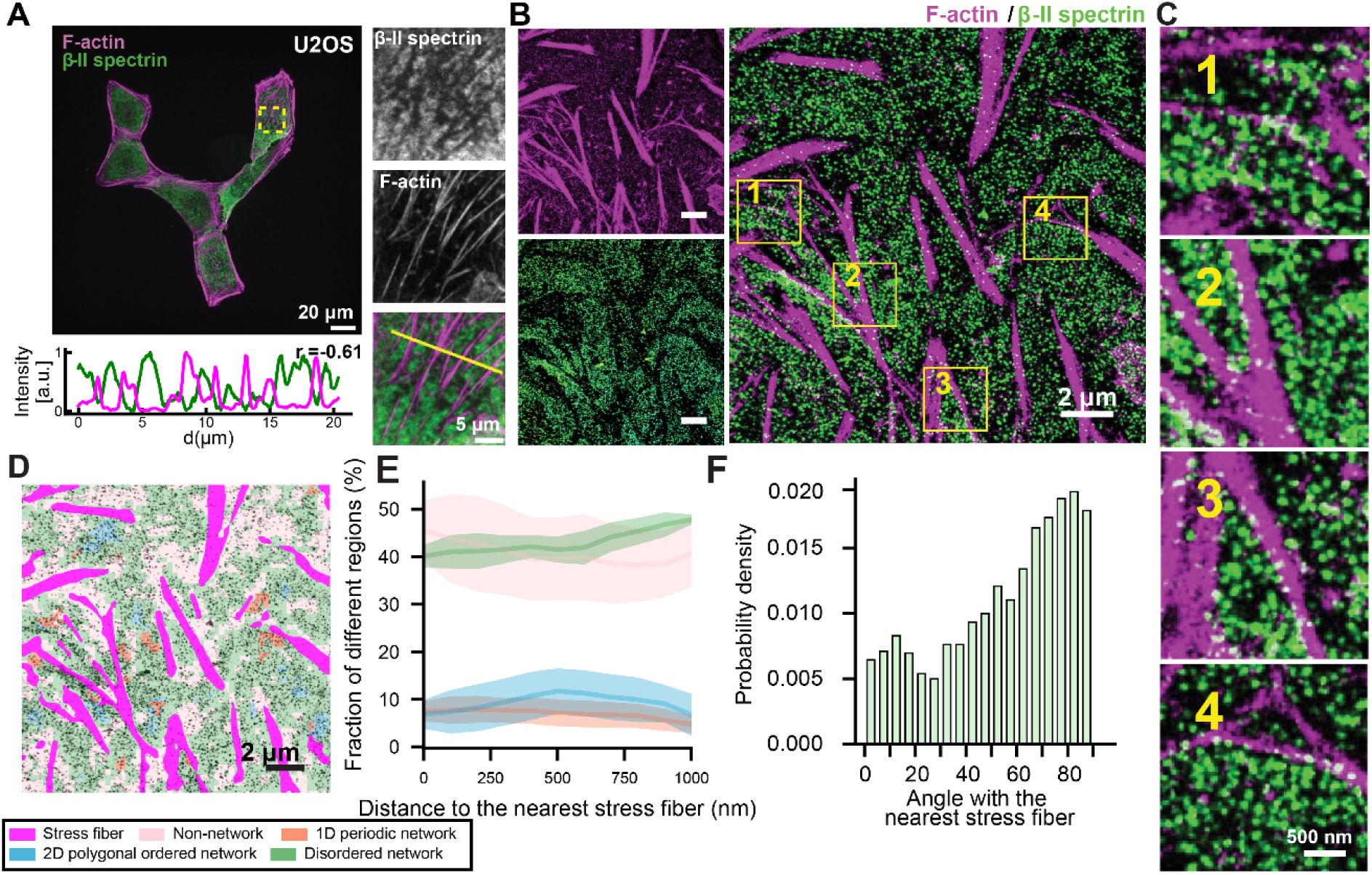
The MPS preferentially forms 1D periodic networks near actin stress fibers, with the periodicity axis oriented perpendicular to the fiber direction. **(A)** Top left: Representative two-color confocal images of U2OS cells co-stained for βII-spectrin (green, βII-spectrin antibody) and F-actin (magenta, phalloidin). Scale bar: 20 µm. Right: Magnified view of the boxed region. Bottom left: Fluorescence intensity profiles of βII-spectrin (green) and F-actin (magenta) along the yellow line showing anticorrelation (Pearson correlation coefficient *r* = - 0.61). Scale bar: 20 µm. **(B)** SMLM (STORM) image of βII-spectrin (green) in a U2OS cell, overlaid with aconfocal image of F-actin (magenta) from the same field of view. Scale bars: 2 µm. **(C)** Magnified views of the four boxed regions shown in (B). Scale bars: 500 nm. **(D)** CLaSSiNet segmentation of the same cell shown in (B), classifying βII-spectrin organization into four MPS structural states: 1D periodic network, 2D ordered network, disordered network, and non-network regions. Scale bars: 2 µm. **(E)** Cell area fractions in the SMLM images exhibiting the four different MPS structural states, plotted as a function of their distance to the nearest actin stress fibers. **(F)** Violin plots of 1D regularity score for cellular regions with their distance to the nearest actin stress fibers, with the median shown as the center line. *p*-values were calculated using a two-sided unpaired Student’s *t*-test. **(G)** Distribution of the relative orientation angles between actin stress fibers and the 1D periodicity axis of adjacent MPS regions, revealing preferential perpendicular alignment.

### 2.7 Comparative Analysis Reveals Cell-Type-Specific MPS Organizational Signatures

Finally, we systematically compared the fractions of the four MPS organizational states across neurons, U2OS, and 3T3 cells (**Figure 8A–C**). Neuronal neurites exhibited the highest fraction of 1D periodic network regions, followed by U2OS cells, 3T3 cells, and neuronal somas (**Figure 8D**). In contrast, neuronal somas contained the largest fraction of 2D polygonal ordered regions, followed by U2OS cells, neuronal neurites, and 3T3 cells (**Figure 8E**). Disordered network regions showed relatively similar fractions across neuronal somas, neurites, U2OS, and 3T3 cells (**Figure 8F**). The average 1D period of the periodic network also varied by cell type: neuronal neurites (∼186 nm) > neuronal somas (∼183 nm) > 3T3 cells (∼182 nm) > U2OS cells (∼179 nm) (**Figure 8G**). These compositional and geometric differences were further reflected in the overall 1D and 2D regularity score distributions for each cell type (**Figure S7A,B**).

**Figure 8.**
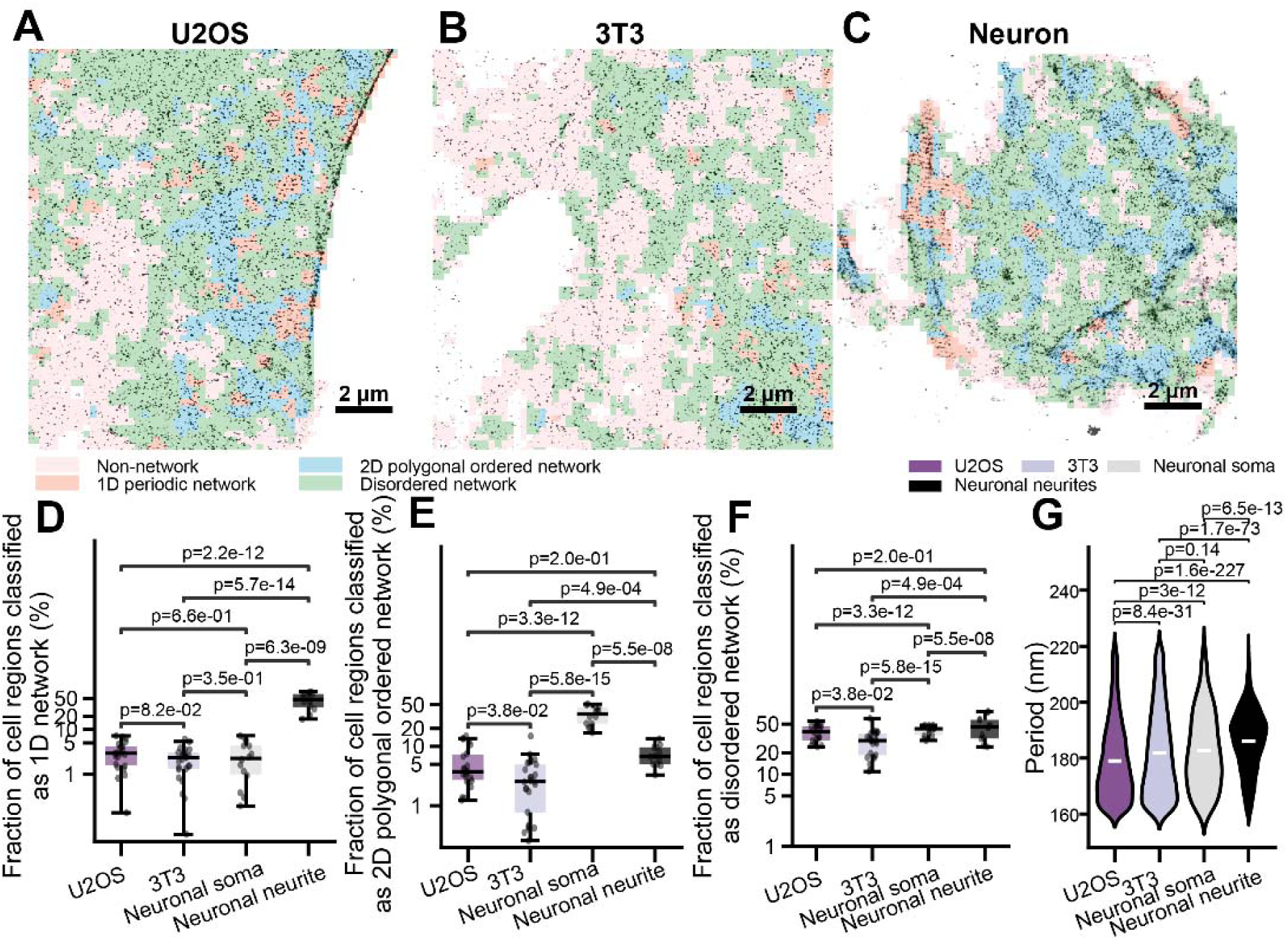
Comparative analysis reveals cell-type-specific MPS architectural signatures. **(A-C)** Representative SMLM (STORM) images of βII-spectrin in a cultured hippocampal neuron (A), a U2OS cell (B), and a 3T3 cell (C), overlaid with CLaSSiNet segmentation into four MPS organizational states: 1D periodic network, 2D polygonal ordered network, disordered network, and non-network regions. Scale bars: 2 µm. **(D, E)** Boxplots showing the cell area fractions exhibiting 1D periodic networks (D) or 2D polygonal ordered network (E) across neuronal somas, neuronal neurites, U2OS cells, and 3T3 cells. **(F)** Boxplots of 1D periodicity periods measured from 1D periodic network regions across neuronal somas, neuronal neurites, U2OS cells, and 3T3 cells. Boxplots show the median and interquartile range (first and third quartiles); whiskers denote the minimum and maximum values excluding outliers. Boxplots show the median and interquartile range (first and third quartiles); whiskers denote the minimum and maximum values excluding outliers. **(G)** Violin plots comparing the first, second, and third nearest-neighbor distance (NND) distributions of 2D polygonal ordered network regions across neuronal somas, neuronal neurites, U2OS cells, and 3T3 cells, with the median shown as the center line. *p*-values were calculated using a two-sided unpaired Student’s *t*-test.

## Discussion

In this work, we developed CLaSSiNet (Classifier of Super-resolution Structural Networks), a modular and quantitative computational pipeline for segmenting and classifying network-like architectures with high classification resolution (256 nm) in super-resolution fluorescence microscopy images. By combining three classifier modules, including connectivity, 1D periodicity, and 2D regularity, we established a framework capable of distinguishing four major organizational states: 1D periodic networks, 2D polygonal ordered networks, disordered networks, and non-network regions. This approach addresses a central challenge in analyzing single-molecule localization microscopy (SMLM) data: although antigens typically appear as sparse localization points rather than complete networks, CLaSSiNet enables reconstruction and classification of the underlying network organization with high classification resolution and robustness to experimental noise.

A key contribution of this work is the systematic benchmarking of network reconstruction and classification methods under both simulated and experimental conditions. By using ground-truth simulated datasets, we demonstrated that our optimized pipelines, particularly Delaunay-based reconstruction combined with triangle area sum, reliably distinguish network from non-network distributions, and that our integrated 1D and 2D classifiers accurately capture structural regularities. Importantly, all three modules remained robust under a range of realistic imaging artifacts, including localization errors, false-positive localizations, and missed detections, underscoring their utility for analyzing diverse SMLM datasets.

Another significant advance of CLaSSiNet is its ability to analyze both link-midpoint (*e.g.*, β-spectrin C-termini) versus node (*e.g.*, actin filaments or β-spectrin N-termini) labeling.. Our analyses revealed that link-midpoint labeling is sufficient to capture connectivity, 1D periodicity and 2D regularity. These findings highlight the generalizability of our framework across different labeling strategies, while also cautioning that reconstructing node distributions from midpoint data introduces additional errors and performs worse than directly analyzing the link-midpoint distributions.

Applying CLaSSiNet to experimental datasets uncovered several new insights into the organization of the membrane-associated periodic skeleton (MPS). In epithelial (U2OS) and fibroblast (3T3) cells, we found that distinct MPS organizational states occupy different subcellular zones, with 1D periodic and 2D ordered regions enriched at cell edges and junctions, while non-network regions dominate cell bodies. Segmented 1D periodic network regions also exhibited larger periods at cell edges and cell–cell junctions compared to the cell body. As previous studies using biophysical techniques and fluorescent biosensors have shown that membrane tension is higher at cell edges^30–32^ and cell-cell junctions^33,34^, it raised the possibility that that local mechanical tension governs MPS architectural state with highly ordered 1D periodic and 2D ordered MPS structures were preferentially formed under high membrane tensions.

We further discover a mechanistic insight that proximity to actin stress fibers influences MPS organization. Specifically, 1D periodic network was more frequently found adjacent to stress fibers, and their periodic axis was preferentially oriented perpendicular to fiber orientation. These results suggest a functional interplay between spectrin and actin-based systems, where the MPS may provide complementary mechanical support and membrane organization in regions flanking contractile actin bundles. Such spatial coordination is likely influenced by local membrane tension, which is known to be elevated near stress fibers^35,36^.

Finally, cross–cell-type comparisons revealed distinct MPS organizational signatures across neuronal somas, neurites, U2OS cells, and 3T3 fibroblasts. Neuronal neurites contained the highest fraction of 1D periodic network regions with the longest average periodicity, whereas neuronal somas exhibited the largest fraction of 2D polygonal ordered networks. U2OS and 3T3 cells displayed intermediate compositions, with 1D periodic fractions lower than in neurites and 2D ordered fractions lower than in neuronal somas. These compositional and geometric differences highlight the adaptability of the spectrin-based MPS, which can assemble into cell-type–specific architectures suited to the distinct structural and functional demands of each cellular context.

Together, these findings establish CLaSSiNet as a versatile and broadly applicable tool for quantitative analysis of nanoscale networks. Beyond the MPS, this framework could be extended to other cytoskeletal, junctional, or organellar networks imaged by SMLM or related super-resolution methods. More broadly, the ability to systematically map nanoscale organization across cell types, subcellular regions, and neighboring structures provides a foundation for uncovering general design principles of cellular networks, as well as their alterations in disease or under mechanical stress. By bridging computational image analysis with cell biology, CLaSSiNet opens new avenues for dissecting how nanoscale architectures contribute to cellular mechanics, signaling, and function.

## Conclusion

In summary, we developed CLaSSiNet, a modular and quantitative computational framework that enables robust segmentation and classification of nanoscale network architectures from super-resolution microscopy data. By integrating connectivity, 1D periodicity, and 2D regularity classifiers, CLaSSiNet accurately distinguishes between distinct structural organizations and remains resilient to experimental noise and labeling variations. Using the MPS network as a model system, we demonstrated that this approach reveals cell-type-specific and subcellular variations in MPS organizations, as well as its mechanical coordination with actin stress fibers, where 1D periodic networks preferentially form near stress fibers and align perpendicularly to their orientation. These findings underscore the adaptability of spectrin-based scaffolds to local mechanical environments and establish CLaSSiNet as a broadly applicable tool for quantitative mapping of nanoscale architectures. More broadly, this framework lays the groundwork for elucidating general design principles that govern the assembly, organization, and function of cytoskeletal, junctional, and organellar networks across diverse biological systems.

## Methods and Materials

### Antibodies

The following primary antibodies were used in this study: mouse anti-βII spectrin antibody 1:200 dilution for IF (Santa Cruz, sc-515592), mouse anti-βII spectrin antibody 1:200 dilution for IF (BD Biosciences, 612563), rabbit anti-adducin antibody 1:500 dilution for IF (Abcam, ab51130), goat anti-βIII spectrin antibody 1:100 dilution for IF (Santa Cruz, sc-9660), mouse anti-βIII spectrin antibody 1:100 dilution for IF (Santa Cruz, sc-515737) The following secondary antibodies were used in this study: Alexa Fluor-647-conjugated donkey anti-rabbit IgG antibody 1:800 for IF (Jackson ImmunoResearch, 711-605-152), Alexa Fluor-647-conjugated donkey anti-mouse IgG antibody 1:800 for IF (Jackson ImmunoResearch, 715-605-151), Alexa Fluor-647-conjugated donkey anti-goat IgG antibody 1:800 for IF (Jackson ImmunoResearch, 705-605-147), CF583R-conjugated secondary antibodies were made by conjugating the unlabeled secondary antibodies with CF583R succinimidyl ester (Biotium, 96084) to achieve a labeling efficiency of ∼2 dyes per antibody, as previously described.

### Cell culture

U2OS and 3T3 cells were grown in DMEM (Gibco; 11966025) supplemented with 2 mM L-glutamine (Gibco; 25030081), 10% fetal bovine serum (Fisher Scientific; SH3008802HI), and 1% penicillin/streptomycin (Gibco; 15140122). Cultures were maintained in a humidified incubator at 37 °C with 5% CO_2_. The cells were plated on precleaned 18-mm coverslips with 10-20% confluency. After 24-48 hours, the cells were fixed and permeabilized for immunostaining.

All animal procedures were performed in compliance with the National Institutes of Health Guide for the Care and Use of Laboratory Animals and were approved by the Pennsylvania State University IACUC. Primary cultures of hippocampal neurons were prepared as previously described^16,20,23^. Briefly, timed-pregnant CD-1 IGS mice (Charles River Laboratories, Wilmington, MA; 022) were euthanized, and hippocampi were isolated from E18 mouse embryos. The dissected tissues were enzymatically dissociated with 0.25% trypsin-EDTA (1x) (Sigma, T4549) at 37 °C for 15 minutes. Following digestion, the hippocampal tissues were washed three times with Hanks’ Balanced Salt Solution (HBSS) (Thermo Fisher Scientific, 14175095), and then transferred to NbActiv1 culture medium (Transnetyx, NB1500 ml), a pre-mixed formulation comprising Neurobasal, B27, and Glutamax, with the addition of 100 μg/ml Primocin (InvivoGen, ant-pm-2). The tissues were gently triturated in the culture medium until a single-cell suspension was achieved, ensuring no tissue clumps remained. Dissociated cells were then counted and plated onto poly-D-lysine-coated 18-mm coverslips (Neuvitro, GG-18-1.5-PDL). Cultures were maintained in a humidified incubator at 37 °C with 5% CO. Half of the medium volume was replaced every five days to maintain optimal culture conditions. Neurons were fixed between 21 and 28 days in vitro for subsequent experiments described in this study.

### Fixation and fluorescence labeling of cells

Cells were fixed with 4% (w/v) PFA and 4% sucrose in PBS for 15-30 min. The fixed cells were blocked with the blocking buffer comprising of 3% (w/v) BSA (Fisher Bioreagents; 9048468), 0.5% (v/v) Triton X-100 (Sigma Aldrich; X100) in PBS (Thermo Scientific; J61196.AP) for 20 min and then incubated with the primary antibodies in the blocking buffer overnight at 4°C, followed by three washes with PBS. The cells were further incubated with the corresponding dye-conjugated secondary antibodies for 60 min at room temperature (RT), washed thoroughly with PBS four times at RT, and then stored at 4°C. To label actin stress fibers, cells were incubated with CF583R- or AF647-conjugated phalloidin (500 nM in PBS; Biotium, 00064; Thermo Fisher, A22287) either overnight at 4 °C or for 1–4 h at room temperature, followed by three washes with PBS.

### STORM imaging

The STORM setup was built on a Nikon Eclipse-Ti2 inverted microscope, equipped with a 100x (N.A. 1.45) Plan Apo oil-immersion objective (Olympus) and an EM-CCD camera (Andor iXon Life 897). 405-nm (Coherent, OBIS 405 nm LX, 1265577; 140 mW), 488-nm (Coherent, Sapphire 488-500 LPX CDRH, 1416094; 300 mW), 560-nm (MPB Communications Inc., 2RU-VFL-P-1500-560-B1R; 1500 mW) and 642-nm (MPB Communications Inc., 2RU-VFL-P-2000-642-B1R; 2000 mW) lasers were introduced through the back port of the microscope. The laser beams were shifted toward the edge of the objective to achieve near-total internal reflection (TIR) illumination, ensuring selective excitation of fluorophores within a few micrometers of the coverslip surface. For single-color 3D STORM imaging, AF647 was imaged. A cylindrical lens was inserted into the detection path to introduce ellipticity into the point spread function, allowing determination of the *z*-positions of single molecules from the ellipticity of their images. Samples were imaged in an oxygen scavenging imaging buffer, containing 100 mM Tris-HCl (pH 7.5), 100 mM cysteamine (Sigma-Aldrich, G8270-100G), 5% (w/v) glucose (Sigma, G8270-100G), 0.8 mg/mL glucose oxidase (Sigma-Aldrich, G2133-10KU), and 40 µg/mL catalase (Sigma-Aldrich, C100-50MG). During imaging, continuous illumination of 642-nm or 560-nm laser (∼2 kW/cm^2^) was used to excite AF647 or CF583R molecules, respectively, switching them into the dark state. Continuous illumination of the 405-nm laser (0-1 W/cm^2^) was used to reactivate the fluorophores to the emitting state, and the illumination power was controlled so that at any given time, only a small, optically resolvable fraction of the fluorophores in the field of view were in the emitting state. A typical single-color 3D STORM image was reconstructed from ∼30,000 image frames acquired at a frame rate of 110 Hz. Super-resolution images were reconstructed from the molecular coordinates by depicting each location as a 2D Gaussian peak.

### STORM image preprocessing

Raw SMLM data from STORM imaging was processed to generate localization coordinates using Insight3 as previously described^16,20,23^. The rendered STORM images were generated by binning the SMLM localization data into a 2D histogram with a pixel size (bin size) of 16 nm. This value was chosen to be on the order of the estimated localization precision (∼10-20 nm) and to comfortably satisfy the Nyquist sampling theorem for the finest periodic features of interest (∼95 nm), which requires a pixel size smaller than approximately 47.5 nm. The resulting 2D histogram was then used to produce the final rendered STORM image.

To identify individual molecular clusters from SMLM localization data, we implemented a two-pass, coarse-to-fine strategy balancing computational efficiency and precision. In the first pass, Laplacian of Gaussian (LoG) blob detection (scikit-image blob_log) was applied to contrast-enhanced rendered STORM images (16 nm/pixel) to rapidly identify candidate regions of interest likely containing molecular clusters. The algorithm searched across multiple scales using 20 logarithmically-spaced levels with intensity thresholds of 0.1. In the second pass, for each candidate ROI, we extracted the corresponding raw localization coordinates (< 20 nm precision) from the original dataset within a 50 nm search radius. HDBSCAN (Hierarchical Density-Based Spatial Clustering, min_cluster_size=10, min_samples=5) was then applied to this subset to accurately define final cluster membership, automatically discarding outlier localizations and determining precise cluster centers directly from the nanoscale point cloud. This hybrid approach leverages the speed of image-based detection for large fields of view while retaining the nanometer-scale accuracy of point-cloud clustering, avoiding information loss from image rendering.

### SMLM image simulations

To evaluate CLaSSiNet under realistic imaging conditions, we generated synthetic single-molecule localization microscopy (SMLM) datasets that emulate network-like cellular structures at the level of individual molecular detection events. Network elements were represented as localization clusters, where each cluster corresponds to either a node (e.g., a filament junction) or a link-midpoint (e.g., the midpoint of a spectrin tetramer connecting adjacent nodes). Each cluster consisted of multiple simulated localizations distributed within a small radius around its centroid, capturing the spatial spread expected from experimental SMLM imaging.

To ensure realism, key simulation parameters were empirically derived from representative experimental datasets (**Figure S1**). First, we quantified fluorophore blinking statistics by fitting Log-Normal distributions to the experimental measurements of (1) the number of localization events per molecular/localization cluster (**Figure S1A**) and (2) the localization precision of single emitters (**Figure S1B**). These fitted distributions were sampled during simulation to assign the number of localizations and per-localization positional uncertainty for each cluster, thereby reproducing realistic SMLM variability.

Simulated networks were initially constructed as perfect ordered structures: parallel lines with equal spacing (e.g., 190 nm for 1D periodic network^20^ for 1D periodic networks and hexagonal lattices for 2D polygonal ordered networks. We then introduced experimentally motivated perturbations, including a missed-detection rate, a false-positive localization rate, and lateral positional deviations of cluster centroids. These perturbations were systematically tuned so that the simulated datasets reproduced the experimentally observed distributions of nearest-neighbor distances (NNDs) for each organizational state (**Figure S1C, D**). Additionally, the average spacing between adjacent cluster centroids perpendicular to the 1D periodicity axis was measured from experimental data (**Figure S1E,F**) and used to set the geometric scale of simulated 1D periodic network. Finally, non-network (random) organization was produced using a Completely Spatially Random (CSR) Poisson process, in which localization clusters were placed independently and uniformly within a defined area to match empirically measured localization cluster densities. All simulations used this set of empirically constrained parameters and geometry rules, and thus provide a controlled yet biologically faithful basis for method validation and benchmarking.

### Generation of composite simulated SMLM images

To assess segmentation performance at interfaces between distinct network organizations, we generated composite SMLM images by combining localization clusters with different structural patterns across irregular boundaries. These boundaries were defined using a multi-step procedure: random noise generation, Gaussian blurring to establish the spatial scale of features, thresholding, and morphological refinement to create smooth, non-linear masks. Each mask delineated two complementary regions within the image, which were then populated with localization clusters representing different organizational states (*e.g.*, 1D periodic network versus random distribution, or 2D polyclonal ordered network versus random distribution) at matched cluster densities. This approach produced composite datasets with known ground-truth spatial domains, enabling quantitative benchmarking of segmentation accuracy across domain interfaces.

### Reconstruction of node positions based on link-midpoint positions

To evaluate how node positions (*e.g.*, labeled at β-spectrin N-terminus for the MPS network) in a biological node-link network can be inferred from SMLM images of link midpoints (*e.g.*, labeled at β-spectrin C-terminus for the MPS network), we developed and tested an algorithm that computationally reconstructs node positions using only the centroid coordinates of link-midpoint localization clusters. The algorithm proceeds in several steps (**Figure S5**). First, initial candidate node positions were identified as vertices of a Voronoi tessellation constructed from the link-midpoint centroids, because these vertices, similar to biological nodes, are equidistant from neighboring link-midpoint clusters. Candidate nodes were then geometrically filtered, retaining only those whose distances to surrounding link-midpoint centroids fell within a physically plausible range. If the internodal distance was smaller than the effective image resolution, the corresponding candidate nodes were consolidated into a single representative node. Finally, an iterative network-assembly step refined the predicted node set: starting from the highest-confidence nodes (*i.e.*, those connected to multiple link midpoints), the algorithm enforced the constraint that each link midpoint connects exactly two nodes. Missing N-terminal partners were computationally added where geometrically feasible to complete N–C–N spectrin tetramer structures. This workflow produced a final predicted map of N-terminal node positions, which was subsequently used for downstream network reconstruction and validation across 1D periodic, 2D polygonal ordered, and disordered network patterns.

### Cellular area determination in STORM images

Cell boundaries were identified from rendered STORM images using fluorescence-intensity–based segmentation. The rendered SMLM image was first divided into a grid with a pixel pitch of 256 × 256 nm. A sliding analysis window (4 × 4 pixels; step = 1 pixel) was then applied to scan the entire image, computing the total fluorescence intensity within each window to generate an intensity heatmap. Pixels with summed intensities below a defined threshold (< 10 arbitrary units) were excluded from subsequent classifier analyses to eliminate regions dominated by background signal. Contour detection was then applied to the retained regions, and only continuous areas exceeding a minimal size (∼5000 nm²) were preserved to further remove noise and spurious detections. This procedure ensured that downstream analyses were restricted to high-confidence cellular areas. For neurite–soma segmentation, morphological opening was applied to the cell masks to selectively remove thin neurites while preserving the larger soma region.

### Connectivity Classifier Module

To classify local regions as network or non-network from SMLM data, we developed a connectivity classifier module that reconstructs node–link organization from localization cluster centroids and quantifies connectivity within spatial windows. Two reconstruction strategies were implemented: range-based and Delaunay-based. In the range-based method, centroid pairs whose separations fell within biologically constrained distances (160–220 nm for node images and 40–220 nm for link-midpoint images) were connected as putative spectrin links; intersecting links were resolved by randomly removing one. In the Delaunay-based method, a Delaunay triangulation was generated from the centroids, and links (*i.e.*, triangle edges) outside the same distance windows were removed, except when they were the sole out-of-range link in a triangle whose remaining two links were in range, preserving characteristic triangular motifs.

Connectivity was quantified using three metrics normalized per unit area: (1) link count, (2) triangle count, and (3) triangle area sum. These metrics capture pairwise connectivity, closed-network motifs, and geometric extent, producing six reconstruction–quantification pipelines (2 reconstruction methods × 3 metrics), among which, Delaunay-based method combined with triangle area sum is designated as the optimized connectivity score.

To resolve spatial heterogeneity, a sliding analysis window (1024 nm × 1024 nm; step = 256 nm) was applied to the rendered SMLM image: each window was independently reconstructed and quantified, and the resulting values were mapped back to generate a connectivity score heatmap across the cell. To identify genuine network domains rather than density-driven artifacts, connectivity values in each window were assessed against density-matched null models. For every window, 100 simulated random centroid distributions were generated at the same centroid density, and a connectivity threshold was defined as the 95th percentile of the simulated distribution. Sliding windows with connectivity scores exceeding this density-dependent threshold were classified as significantly connected (5% false-positive rate; *p* < 0.05), enabling robust detection of MPS network regions.

### 1D Network Classifier Module

To identify 1D periodic network regions, each SMLM image was divided into grids using a moving window (1024 × 1024 nm; step = 256 nm). Within each window, 1D periodic network was detected using Fourier-based spectral analysis. Each window was preprocessed by normalization, multiplication with a 2D Hann window to minimize spectral leakage, and zero-padding, followed by a 2D Fast Fourier Transform (FFT) to the frequency domain. The dominant frequency peaks were isolated using an annular mask corresponding to real-space periods of 160–220 nm, and for each window, the orientation (θ) and period (*d*) were extracted using the inverse FFT which generated a grating image representing the best-fitting orientation and period of the candidate 1D network.

Five candidate 1D regularity scores were computed for each window to quantify the degree of 1D periodic order: (1) FFT peak amplitude – the amplitude of the two dominant frequency peaks in the 2D FFT image. (2) 2D Pearson correlation – correlation between the raw SMLM image and the inverse FFT-generated grating. (3) 1D projected Pearson correlation – correlation between 1D profiles of the raw and grating images projected along the grating axis. (4) Localized 2D Pearson correlation – correlation between the raw SMLM image and a cropped, elongated grating image to capture local 1D network subregions. (5) Integrated 1D regularity score – a combined metric of (2) and (4) capturing both extended and localized 1D network regions. Among the five scores, integrated 1D regularity score is designated as the optimized 1D regularity score.

For each moving window, the optimized 1D regularity score was computed, and an initial threshold (> 0.6) corresponding to a 1% false-positive rate (*p* < 0.01) was applied to identify high-confidence “seed” pixels as candidate 1D periodic network regions. These seed regions were then expanded via a spatially coherent region-growing algorithm: neighboring pixels were annexed iteratively if their score exceeded a secondary threshold (> 0.45) corresponding to a 5% false-positive rate (*p* < 0.05) and their orientation and period values were consistent with the seed (Δθ < 15°). Post-filtering removed segmented regions below a predefined size, unifying fragmented domains and eliminating spurious pixels. Finally, the refined segmentation produced binary maps distinguishing 1D periodic network from non-1D regions.

### 2D Network Classifier Module

To quantify 2D polygonal network regularity, we applied the same moving-window approach (1024 × 1024 nm; step = 256 nm), partitioning the SMLM images into local grids. For each window, we calculated four candidate 2D regularity scores based on three geometric features derived from the reconstructed network using the Connectivity Classifier Module: (1) Neighbor count similarity – the number of nodes directly connected to a given node. The 2D regularity score is defined as the inverse of the deviation in neighbor count between a node and its immediate neighbors. (2) Link-length similarity – the distribution of lengths of links connecting a node to its neighbors. The score is the inverse of the deviation in link-length distributions between a node and its neighbors. (3) Link-angle similarity – the distribution of angles formed between links connected to a node. The score is the inverse of the deviation in angular distributions between a node and its neighbors. (4) Integrated 2D regularity score – a composite metric combining all three geometric features to capture overall local network regularity. To optimize the combination of these features for distinguishing ordered from disordered networks, we trained a Linear Discriminant Analysis (LDA) model using simulated ground-truth datasets containing 2D ordered and disordered networks. For each moving window, the 2D regularity score of the window was calculated as the average score of all nodes contained within it, generating a heatmap representing the local degree of 2D regularity. This approach enables robust identification and mapping of polygonally ordered network regions across the cellular area. Among the four scores, integrated 2D regularity score is designated as the optimized 2D regularity score.

### CLaSSiNet workflow

We integrated the three classifier modules into a unified analysis pipeline, CLaSSiNet, to segment SMLM node images into four categories of cellular regions corresponding to four distinct network organizational states: 1D periodic network, 2D polygonal ordered network, disordered network, and non-network. The workflow proceeds sequentially as follows: (1) Connectivity Classifier Module – segments the SMLM image into network and non-network regions; (2) 1D Network Classifier Module – identifies 1D periodic networks within the network regions; (3) 2D Network Classifier Module – classifies the remaining network regions to detect 2D polygonal ordered networks; (4) Residual classification – remaining network regions are designated as disordered networks. This modular workflow allows systematic, hierarchical classification of network structures, generating spatial maps of four distinct organizational states across the cellular area.

### Actin stress fiber detection

Stress fibers were identified from conventional fluorescence images of F-actin using a dual-threshold segmentation approach. Images were normalized (0-255) and Gaussian-blurred (5×5 kernel) to reduce noise, then segmented using combined global (intensity > 20) and local adaptive thresholding (111-pixel Gaussian window, offset=2). Morphological closing (3×3 kernel) followed by opening (7×7 kernel) removed small gaps and noise. Connected regions were filtered to retain only elongated structures characteristic of stress fibers: eccentricity > 0.8, area > 250um^2^, and major axis length ≥ 2um. For subsequent angle-related analysis, the precise orientation of each identified fiber was determined and manually verified using ImageJ.

### Confocal image cell segmentation and compartment classification

The overall cell boundaries and nuclei were established to define the cellular region. The nuclei were segmented using the simple thresholding of the Hoechst stain. Whole cells were segmented using the watershed algorithm on membrane-stained images. Protein-enriched regions were robustly detected by combining adaptive local thresholding with global top-percentile filtering. This approach targets areas where protein signals are locally high while also being among the brightest global features. The resulting binary masks were further refined by morphological operations and size/intensity filtering (retaining regions >1000 pixels with an average intensity >60% of the maximum). This rigorous process allowed us to assign every pixel to one of five mutually exclusive compartments: blank region, cell body, cell edge, cell-cell junction, and nucleus.

## Acknowledgements

We thank all the members in Prof. Zhou’s laboratory for insightful scientific discussion.

## Author contributions

R.Z. conceived the project. Y.T. and R.Z. designed experiments and interpreted the data. Y.T. performed experiments, designed the algorithm and analyzed the data. R.Z. wrote the manuscript with the input from Y.T. R.Z. acquired funding and supervised the project.

## Consent to Participate

The authors declare their agreement to participate.

## Consent for publication

All the authors declare their agreement to publish.

## Data availability

The data supporting the findings of this study are provided within the paper and its Supplementary Information and are available from the corresponding author upon reasonable request.

## Supplementary Information

**Figure S1.**
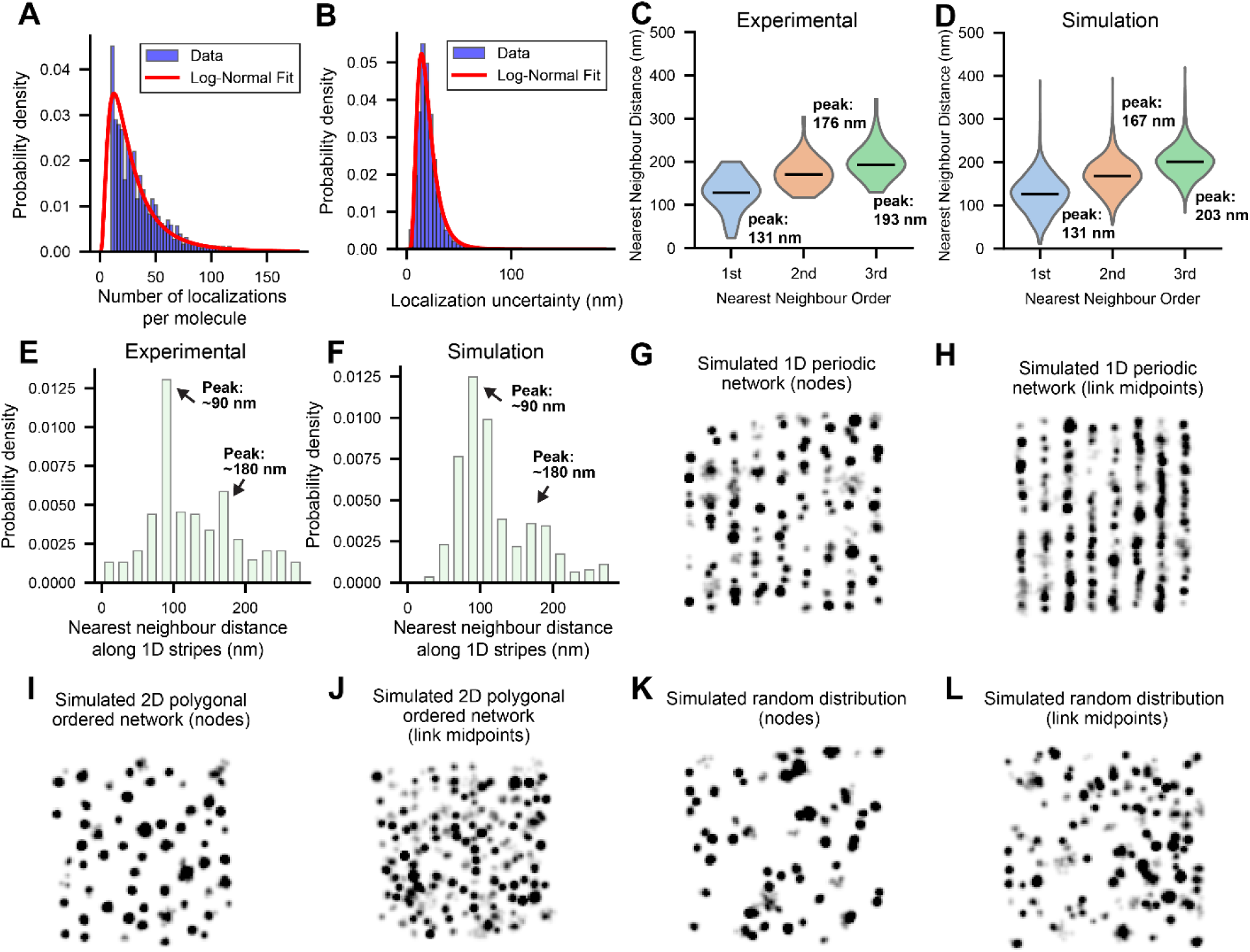
Parameter determination from experimental data for generating simulated SMLM images. **(A, B)** Distributions of the number of single-molecule localizations per molecular cluster (A) and localization uncertainties (i.e., the standard deviation of the Gaussian fit to single emitters) (B) obtained from experimental SMLM images of βIII-spectrin N-termini (labeling actin nodes) in neurons. Red solid lines represent Log-Normal probability density function (PDF) fits. These distributions were used to generate molecular clusters in simulated SMLM images that closely mimic experimental data. In the simulations, molecular clusters correspond to either nodes or the midpoints of links in the node-link network. **(C, D)** Violin plots of nearest-neighbor distance (NND) distributions for the first, second, and third nearest neighbors determined from the 2D polygonal ordered MPS network regions in the experimental SMLM images of actin nodes (i.e., N-terminus of βIII-spectrin immunolabled) (C) and obtained from simulated SMLM node images (D). Peak values of the distributions are indicated. **(E, F)** Distributions of inter-cluster distances perpendicular to the 1D periodicity axis and of periods along the 1D periodicity axis, determined from the 1D periodic regions of experimental SMLM node images (e) and obtained from simulated SMLM node images (F). **(G-J)** Representative simulated SMLM images generated using the parameters derived from panels (A-F), showing examples of node images (G) and link-midpoint images (H) for 1D periodic network, as well as examples of node images (I) and link-midpoint images (J) for 2D polygonal ordered network. **(K)** Same as (I) but for nodes with random distributions and equal node density as in (I). **(L)** Same as (J) but for link-midpoints with random distributions and equal link-midpoint density as in (J).

**Figure S2.**
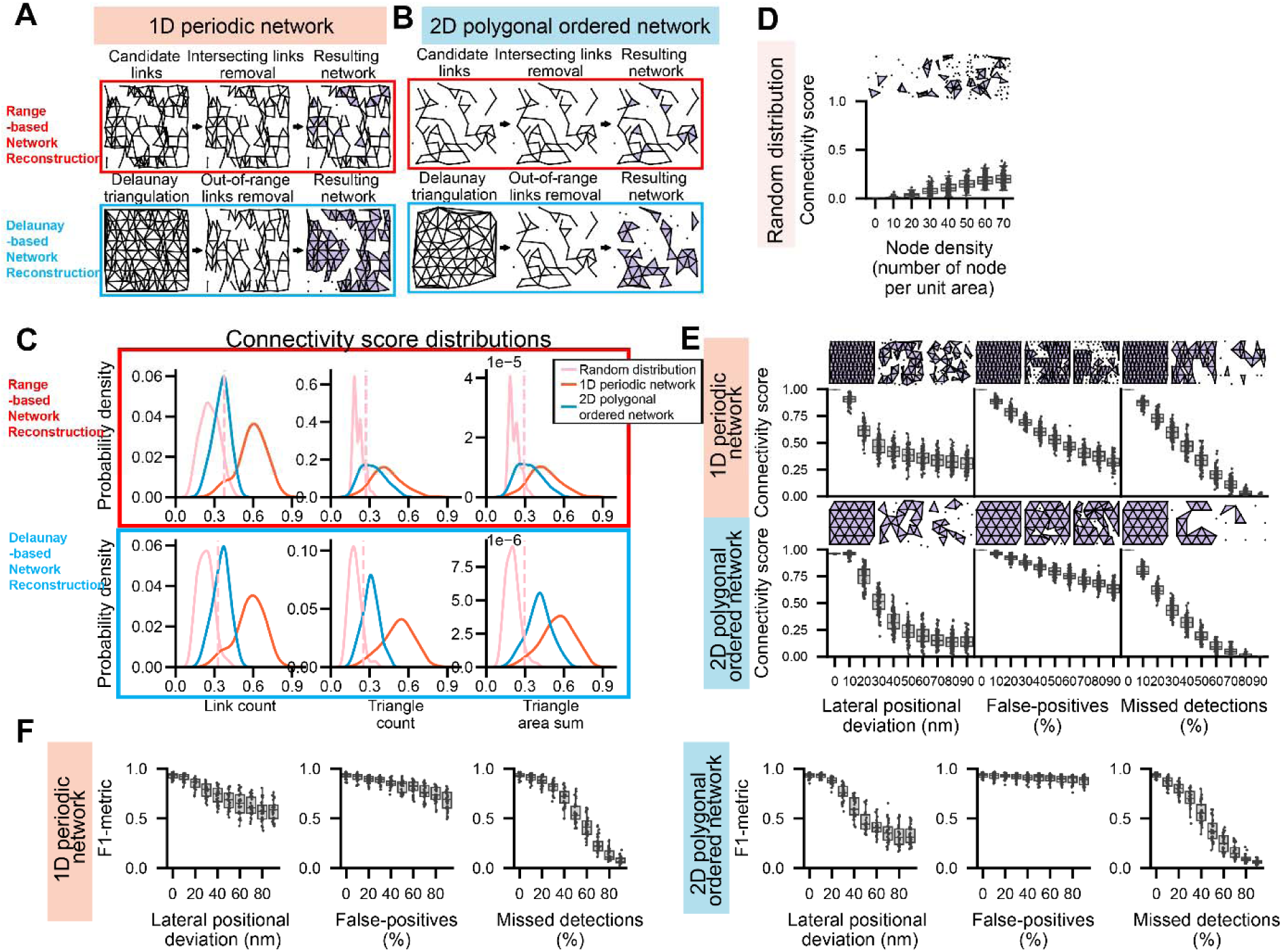
Optimization of the Connectivity Classifier Module. **(A, B)** Steps for range-based (top) and Delaunay-based (bottom) network reconstruction, shown for simulated SMLM node images with 1D periodic (A) and 2D polygonal (B) distributions. **(C)** Kernel density estimation (KDE) plots showing the distributions of link count, triangle count, and triangle area sum calculated from simulated SMLM node images representing random, 1D periodic, and 2D polygonal networks, using range-based (top, in red box) and Delaunay-based (bottom, in blue box) network reconstruction. Dashed lines indicate the 95th percentile of the KDE distribution calculated from simulated SMLM node images with random node distributions, which is used as the threshold to segment network versus non-network regions. **(D)** Boxplots showing that connectivity scores shift to higher values as the node density increases in simulated random node distributions. A node-density-dependent threshold (defined as the 95th percentile of the KDE distribution from simulated random node distributions at each node density) is therefore applied for network segmentation. **(E,F)** Robustness analysis of the optimized Connectivity Classifier Module for quantifying simulated SMLM node images containing 1D periodic, and 2D polygonal networks, under varying added experimental noise, including lateral localization deviation, added false-positive nodes (as a percentage of the total true nodes), and missed detections (as a percentage of the total true nodes). Connectivity scores (E) and F1 metrics (F) under these conditions are shown. Boxplots show the median and interquartile range (first and third quartiles); whiskers denote the minimum and maximum values excluding outliers.

**Figure S3.**
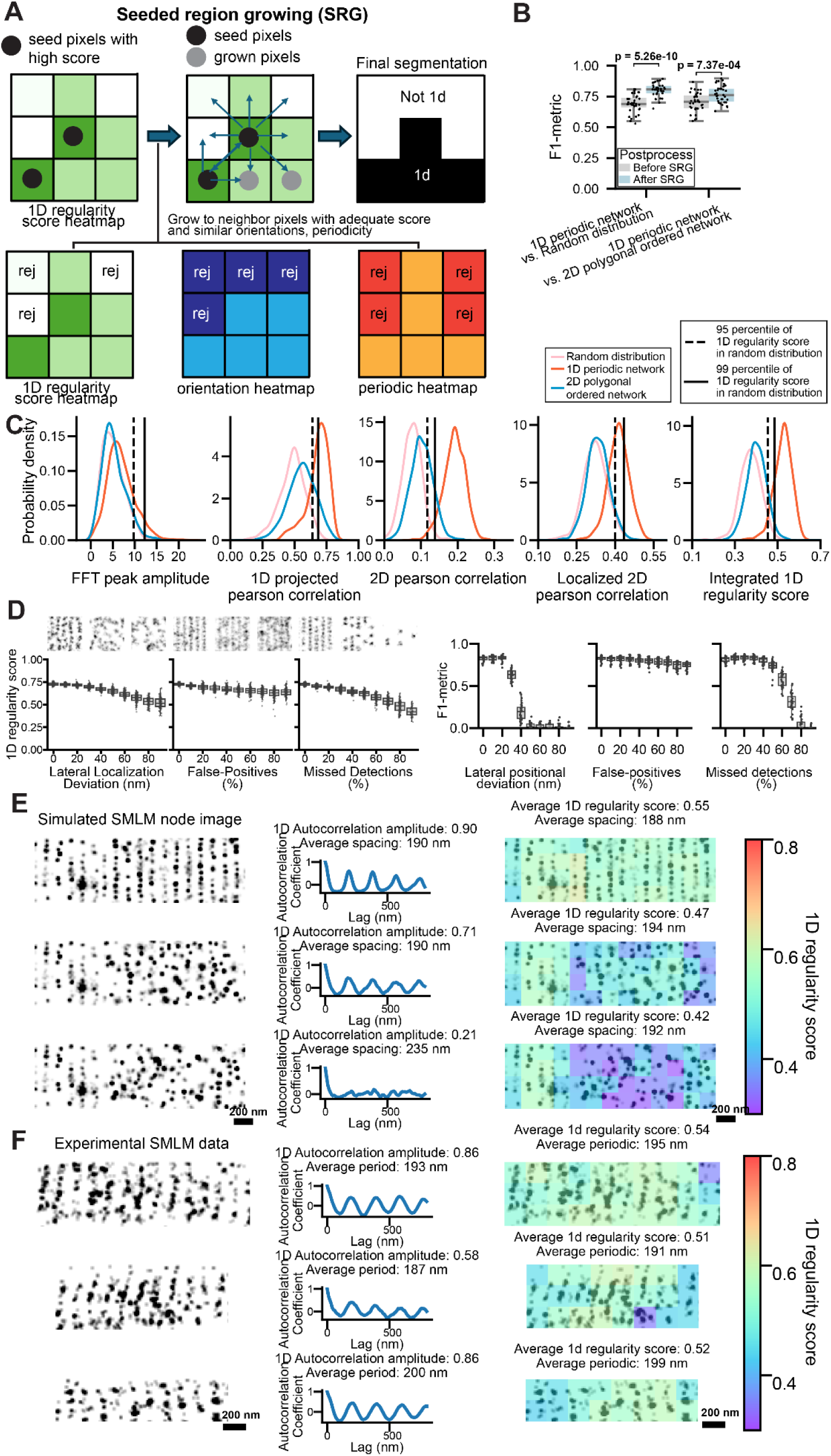
Optimization of the 1D Network Classifier Module. **(A)** Schematic illustrating the segmentation method based on a “seed region growing” (SRG) algorithm. A first threshold (99th percentile) was applied to the 1D regularity score heatmap to identify “seed” pixels with high 1D regularity scores, generating an initial segmentation of 1D network regions. Pixels adjacent to these seeds that exceeded a second threshold of 1D regularity score (95th percentile) were then evaluated based on the orientation and period heatmaps: neighboring pixels were accepted or rejected depending on whether their orientation and period values were close to those of nearby seed pixels, thereby refining the segmentation boundary. **(B)** Boxplots of F1-metrics quantifying segmentation performance in distinguishing 1D periodic network regions from random or 2D polygonal network regions, before and after the “grow” refinement step in the seed-and-grow method. **(C)** Kernel density estimation (KDE) plots showing the distributions of FFT peak amplitude, 2D Pearson correlation, 1D projected Pearson correlation, Localized 2D Pearson correlation, and integrated 1D regularity score, calculated from simulated SMLM node images representing random, 1D periodic, and 2D polygonal networks. Solid black lines and dashed lines indicate the 99th and 95th percentiles of the KDE distribution, respectively, calculated from simulated SMLM node images with random node distributions, which are used as the thresholds to segment 1D periodic network versus non-1D-network regions. **(D)** Robustness analysis of the optimized 1D Network Classifier Module applied to simulated SMLM node images containing 1D periodic networks under varying levels of experimental noise, including lateral localization deviation, added false-positive nodes (as a percentage of the total true nodes), and missed detections (as a percentage of the total true nodes), as well as under different mean inter-cluster distances perpendicular to the 1D periodicity axis. Both connectivity scores (left) and F1 metrics (right) under these conditions are shown. **(E, F)** Comparison between 1D autocorrelation amplitudes determined by traditional 1D autocorrelation analysis and 1D regularity scores determined by the 1D Network Classifier Module, calculated from the same set of simulated SMLM images (E) or experimental SMLM images (F). Scale bars: 200 nm. Boxplots show the median and interquartile range (first and third quartiles); whiskers denote the minimum and maximum values excluding outliers. *p*-values were calculated using a two-sided unpaired Student’s *t*-test.

**Figure S4.**
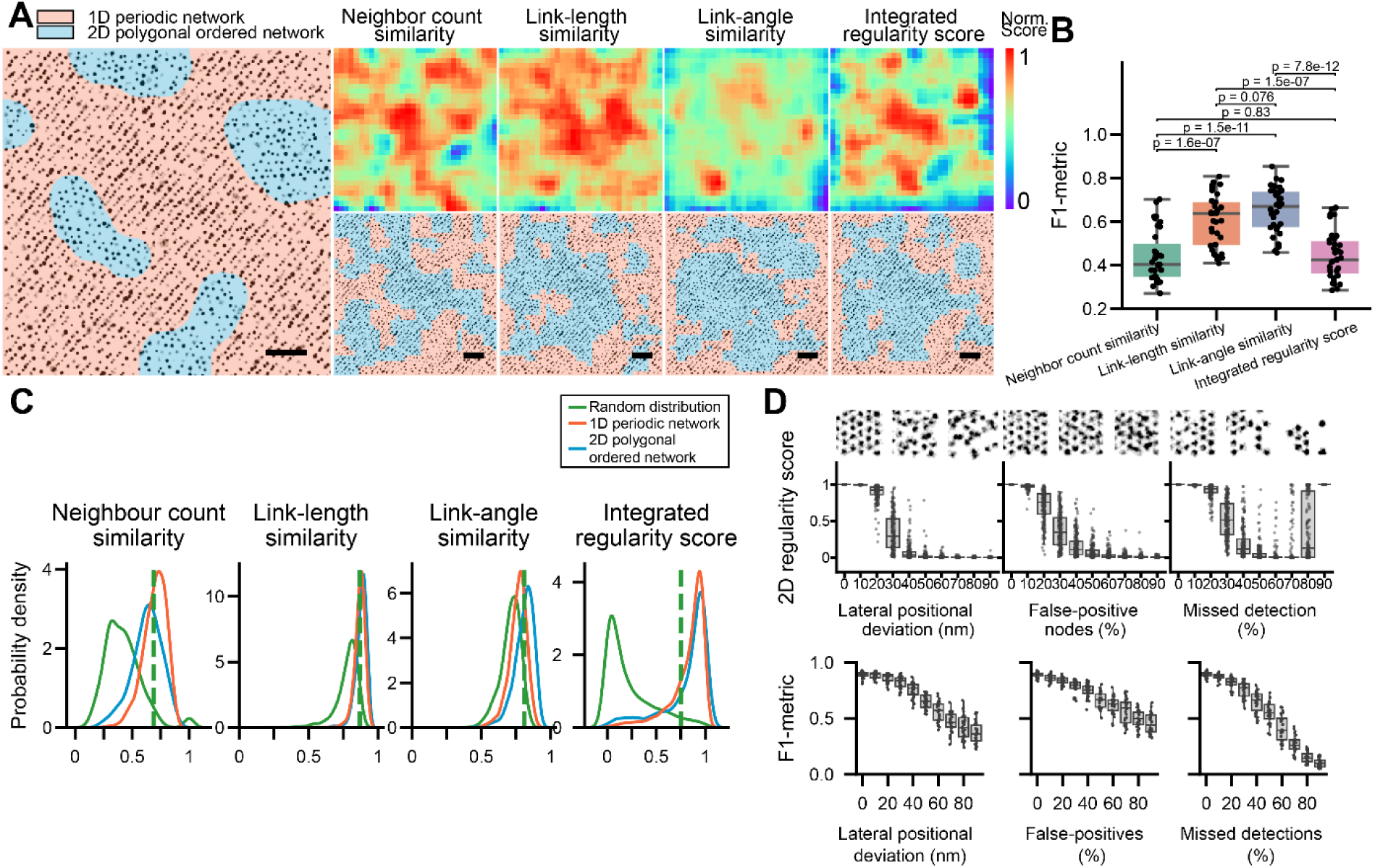
Optimization of the 2D Network Classifier Module. **(A)**Left: Representative simulated SMLM node image containing embedded islands of nodes arranged in 2D polygonal distribution, surrounded by nodes of equal density arranged in a 1D periodic network. Right: heatmaps (top) and segmented images (bottom) generated for the simulated SMLM image on the left, using the four candidate 2D regularity scoring methods. Scale bars: 1 µm. **(B)** Boxplots of F1-metrics quantifying the segmentation performance in distinguishing 2D polygonal network regions from 1D periodic network regions using the four candidate 2D regularity scoring methods. **(C)** Kernel density estimation (KDE) plots showing the distributions of the four candidate 2D regularity scores calculated from simulated SMLM node images representing random, 1D periodic, and 2D polygonal networks. Dashed lines indicate the 95th percentile of the KDE distribution calculated from simulated SMLM node images with random node distributions, which is used to segment 2D polygonal ordered network regions versus random distributions. This demonstrates that the LDA-based 2D regularity score provides the largest separation between the KDE distributions for 2D ordered and random networks, but less so between the KDE distributions for 2D polygonal ordered and 1D periodic ordered networks. **(D)** Robustness analysis of the optimized 2D Network Classifier Module applied to simulated SMLM node images containing 2D polygonal networks under varying levels of experimental noise, including lateral localization deviation, added false-positive nodes (as a percentage of the total true nodes), and missed detections (as a percentage of the total true nodes). Both connectivity scores (left) and F1 metrics (right) under these conditions are shown. Boxplots show the median and interquartile range (first and third quartiles); whiskers denote the minimum and maximum values excluding outliers. *p*-values were calculated using a two-sided unpaired Student’s *t*-test.

**Figure S5.**
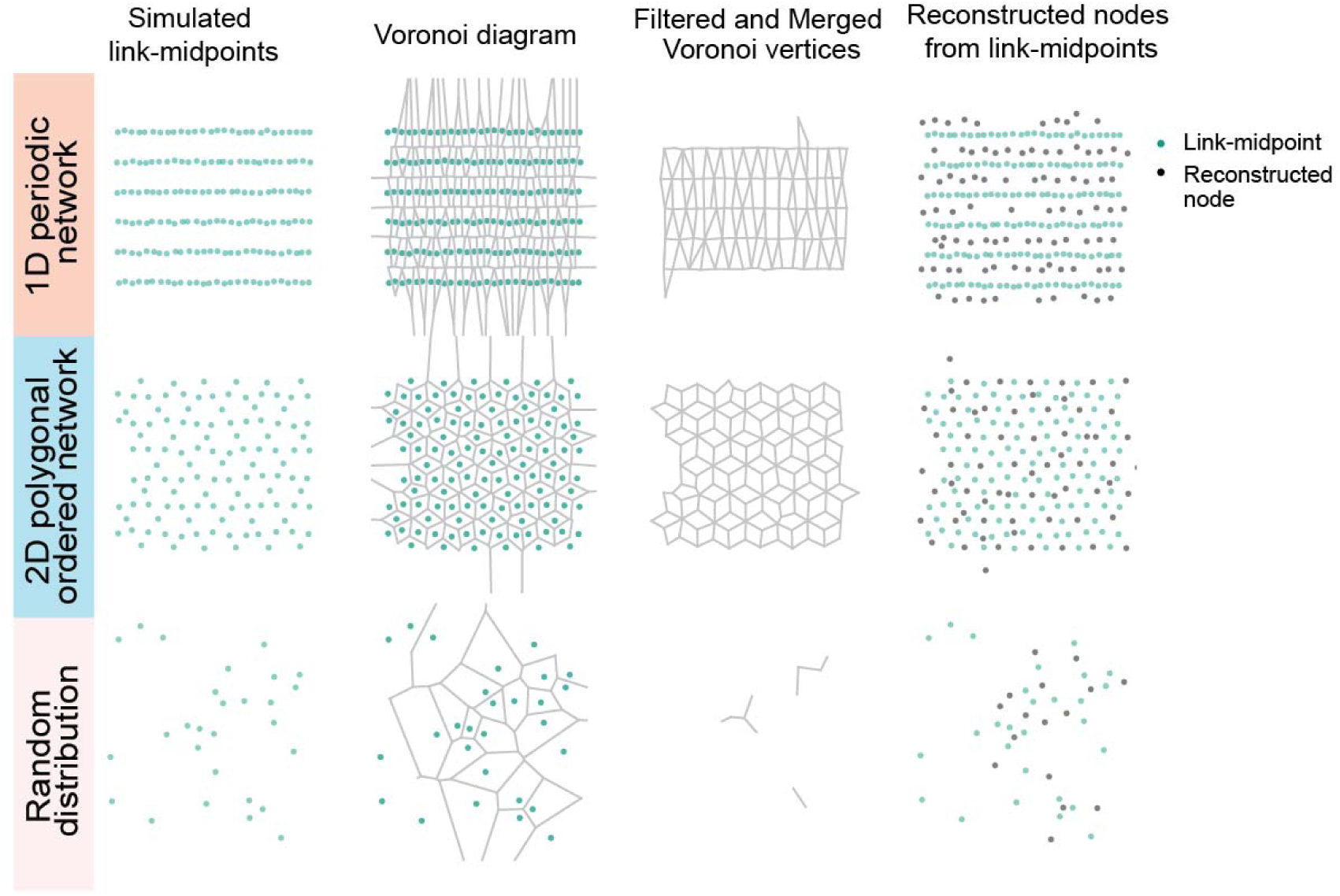
Voronoi tessellation-based algorithm to estimate node locations from link-midpoint distributions. Schematics illustrating the geometric principle of the algorithm designed to reconstruct N-terminal node positions from the simulated SMLM link-midpoint images for 1D periodic network, 2D polygonal ordered network, and random distributions, using Voronoi tessellation-based algorithm.

**Figure S6.**
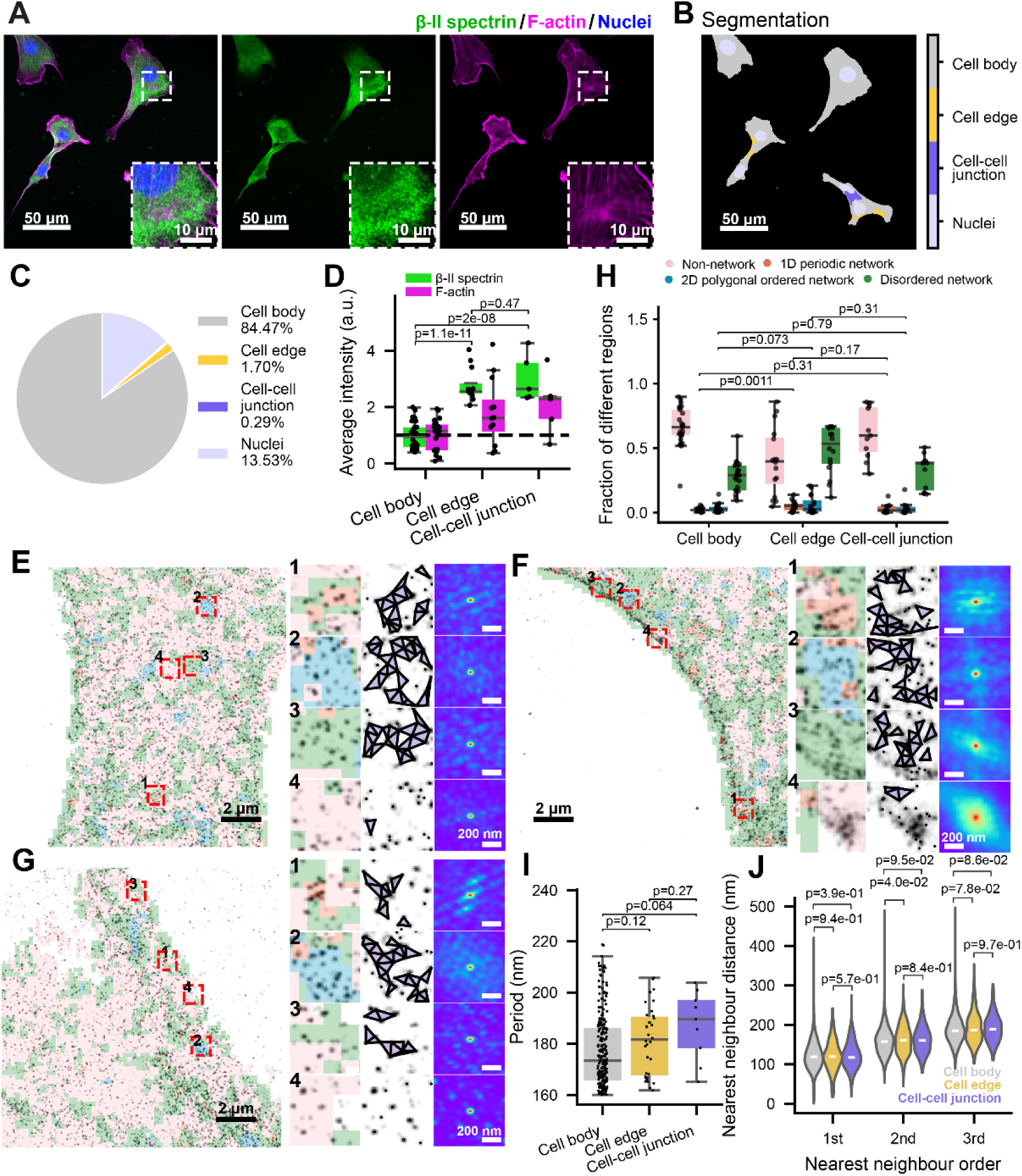
CLaSSiNet reveals distinct MPS organizations across three subcellular zones in fibroblasts. **(A)** Representative three-color confocal images of 3T3 cells co-stained for nuclei (blue, Hoechst), βII-spectrin (green, βII-spectrin antibody), and F-actin (magenta, phalloidin). Scale bar: 50 µm. Inset: Magnified view of the boxed region. Scale bar: 10 µm. **(B)** Segmented image corresponding to (A), showing four subcellular zones: nucleus, cell body, cell edge, and cell–cell junction. Scale bar: 50 µm. **(C)** Quantified area fractions of the four subcellular zones. **(D)** Boxplots showing average fluorescence intensities of βII-spectrin and F-actin across the three non-nuclear subcellular zones (cell body, cell edge, and cell–cell junction). **(E–G)** Left: Representative SMLM (STORM) images of βII-spectrin in three subcellular zones, cell body (e), cell edge (F), and cell–cell junction (G), overlaid with CLaSSiNet segmentation into four MPS organizational states: 1D periodic network, 2D polygonal ordered network, disordered network, and non-network regions. Scale bar: 2 µm. Middle: Magnified views of boxed regions enriched in each of the four organizational states. Right: Reconstructed node-link networks and corresponding 2D autocorrelation maps confirming the expected MPS organizational patterns. Scale bar: 200 nm. **(H)** Boxplots showing the area fractions of the four MPS organizational states across the three subcellular zones. **(I)** Boxplots of 1D periodicity periods measured from 1D periodic network regions across the three subcellular zones. Boxplots show the median and interquartile range (first and third quartiles); whiskers denote the minimum and maximum values excluding outliers. **(J**) Violin plots comparing the first, second, and third nearest-neighbor distance (NND) distributions of 2D polygonal ordered network regions across the three subcellular zones. The center lines indicate the median. *p*-values were calculated using a two-sided unpaired Student’s *t*-test.

**Figure S7.**
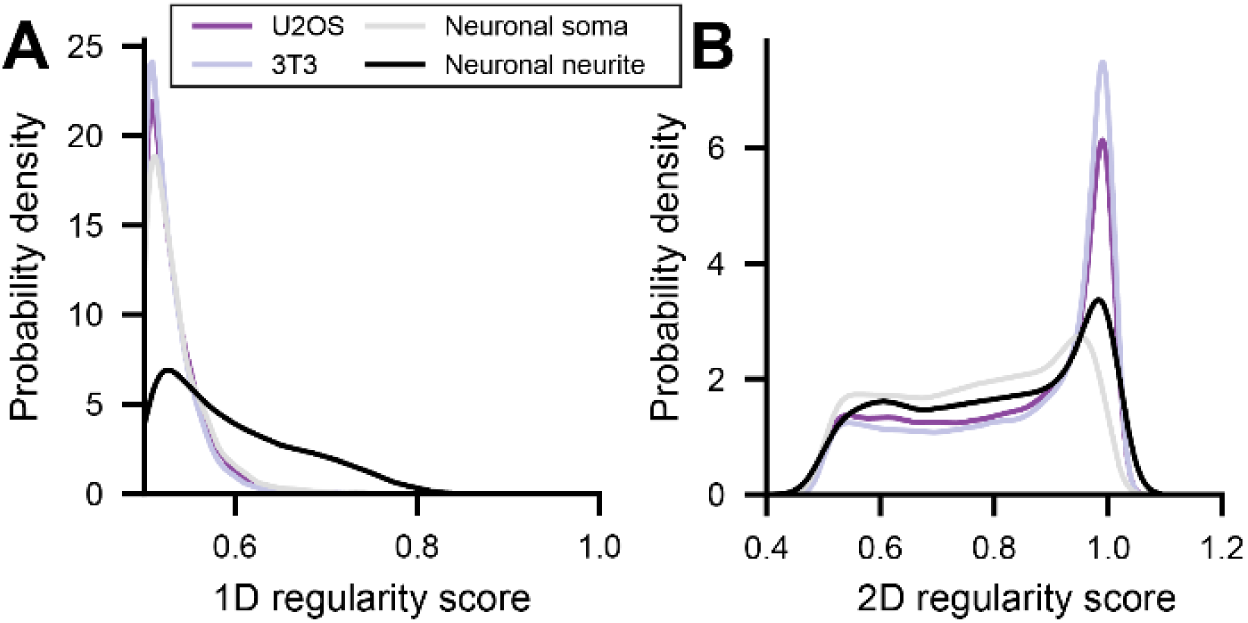
Distributions of 1D and 2D regularity scores for U2OS cells, 3T3 cells, neuronal somas, and neuronal neurites. **(A, B)** Kernel density estimate (KDE) plots showing the distributions of the 1D regularity score (A) and the 2D structural regularity score (B) for all network regions across U2OS cells, 3T3 cells, neuronal somas, neuronal neurites.

